# Machine Learning Reveals Immediate Disruption in Mosquito Flight when exposed to Olyset Nets

**DOI:** 10.1101/2025.02.05.636601

**Authors:** Yasser M. Qureshi, Vitaly Voloshin, Amy Guy, Hilary Ranson, Philip J. McCall, Cathy E. Towers, James A. Covington, David P. Towers

## Abstract

Insecticide-treated nets (ITNs) remain a critical intervention in controlling malaria transmission, yet the behavioural adaptations of mosquitoes in response to these interventions are not fully understood. This study examined the flight behaviour of insecticide-resistant (IR) and insecticide-susceptible (IS) Anopheles gambiae strains around an Olyset net (OL), a permethrin-impregnated ITN, versus an untreated net (UT). Using machine learning (ML) models, we classified mosquito flight trajectories with high accuracy (0.838) and ROC AUC (0.925). Contrary to assumptions that behavioural changes at OL would intensify over time, our findings show an immediate onset of convoluted, erratic flight paths for both IR and IS mosquitoes around the treated net. SHAP analysis identified three key predictive features of OL exposure: frequency of zero-crossings in flight angle change, first quartile of flight angle change, and zero-crossings in horizontal velocity. These suggest disruptive flight patterns, indicating insecticidal irritancy. While IS mosquitoes displayed rapid, disordered trajectories and mostly died within 30 minutes, IR mosquitoes persisted throughout the 2-hour experiments but exhibited similarly disturbed behaviour, suggesting resistance does not fully mitigate disruption. Our findings challenge literature suggesting permethrin’s repellency in solution form, instead supporting an irritant or contact-driven effect when incorporated into net fibres. This study highlights the value of ML-based trajectory analysis for understanding mosquito behaviour, refining ITN configurations and evaluating novel active ingredients aimed at disrupting mosquito flight behaviour. Future work should extend these methods to other ITNs to further illuminate the complex interplay between mosquito behaviour and insecticidal intervention.

## 1 Introduction

Mosquito-borne diseases are among the leading causes of human mortality with over 1 million deaths per annum worldwide and they are a barrier to development in countries where they are endemic (1). Control of mosquitoes with insecticide-based interventions has been a critical part of the global strategy to manage mosquito-borne diseases (2). Pyrethroids are among the most widely used insecticides, delivered historically by indoor residual spraying walls and today primarily as insecticide-treated nets (ITNs), they are highly effective in malaria vector control. However, prolonged, widespread use led to the appearance of resistance which spread rapidly in mosquito populations across Africa following the rollout of ITNs in the first years of the 21st century. In that timeframe, the malaria burden in Africa had been reduced by 50%, most of which was attributed to ITNs (3). By 2015, the pace of progress had declined considerably, and malaria cases have plateaued in recent years (4). An understanding of the mechanisms governing the development of insecticide resistance has led to new and more effective strategies to control resistant mosquitoes, including next generation ITNs. Whilst much is known about the genetic mechanisms of insecticide resistance, far less attention has been paid to the understanding the full range of the impacts of pyrethroid exposure on mosquitoes and malaria transmission (5–7).

The Olyset net, in 2001, was the first ITN to be validated under the WHO Pesticide Evaluation Scheme (WHOPES) and today it is used in over 80 countries, protecting nearly 800 million people (8). Despite widespread pyrethroid resistance Olyset nets can still impact resistant mosquitoes by reducing their willingness to feed and/or reducing their lifespan (9,10). Due to their lower cost, they continue to be used in areas where insecticide resistance levels are low despite the widespread availability of next generation of dual active ingredient nets (11,12).

Recent studies have explored behavioural responses of insecticide resistant mosquitoes on a number of ITNs, including Olyset and some dual-active ingredient nets (10). By tracking various mosquito strains around different human-baited ITNs for 2 hours using an infrared video system, it was concluded that were no major differences in the duration or character of net contact, repellency or irritability of the different nets, or of the consequences of exposure on longevity between pyrethroid resistant and susceptible mosquitoes when exposed to ITNs. The analysis primarily used statistical methods to examine parameters, such as net contact duration, mosquito activity decay, and feeding inhibition. Here, we report on the application of machine learning (ML) models to classify mosquito behaviours based on similar data collected from Olyset and untreated nets. By leveraging explainable AI (XAI) techniques, we have revealed further insights into mosquito movement patterns and on subtle behavioural differences between resistant and susceptible strains that were not detected through traditional statistical methods.

ML models can identify flight characteristics that distinguish groups of mosquitoes. For instance, (13) was able to identify behaviours between sexes of mosquitoes with an average balanced accuracy and ROC AUC score of 0.645 and 0.684 – which represented the first work to use ML models to explore mosquito flight behaviour via trajectories. Similar work (14) was also performed to differentiate between insecticide-susceptible (IS) and insecticide-resistant (IR) be-haviours at an untreated net with the best model returning an average balanced accuracy of 0.743 and a ROC AUC score of 0.813. Furthermore, the model was then explained using SHapley Additive eXplanations (SHAP) values (15) revealed that IR mosquitoes tend to exhibit more directed flight paths with some jerky motion – possibly allowing the mosquito to sample attractants and maintain flight towards the highest concentration of these cues and hence a potential bloodmeal.

In this work, we present the first evidence of the behavioural impact of ITNs on mosquito flight using an established ML approach. The study utilised trajectory data of mosquitoes flying around an untreated baited bednet and a treated bednet (Olyset) in order to identify behaviours resulting from exposure to the insecticide. Features of flight were extracted from the trajectories and provided to a ML model to attempt to distinguish between the two net types. Then, SHAP’s analysis was used to explain the model predictions.

## 2 Methods

### 2.1 Dataset Description

The dataset used was generated within laboratory experiments at the Liverpool School of Tropical Medicine (LSTM), UK (10). Within each experiment, unfed adult female mosquitoes were tracked around a human-baited bednet. In 17 of the experiments, the bednet was untreated (i.e., no presence of insecticide), in 23 of the experiments the bednet was a standard pyrethroid-only net (Olyset NetTM). Olyset nets are made from polyethylene fibres with permethrin (2%) incorporated directly into the material during manufacturing, rather than being surface-coated like conventional ITNs. The manufacturer states that this allows the insecticide to gradually migrate to the surface over time, and that Olyset nets must be activated before first used by heating after which time they will provide effective protection for a few years (16). The nets feature a wide mesh size (4mm x 4mm) designed to maximise ventilation while still preventing mosquito entry. Four Anopheles gambaie mosquito strains were used, two insecticide-susceptible (Kisumu and N’goussu) and two insecticide-resistant (VK7 and Banfora). The Kisumu strain, first collected in Kenya in 1975, and the N’gousso strain, established in 2006 from Cameroon, are both insecticide-susceptible mosquito populations (17). In contrast, the VK7 and Banfora strains, derived from Burkina Faso in 2014 and 2015, exhibit high levels of pyrethroid resistance. This resistance is primarily driven by kdr mutations (L1014F) and metabolic adaptations, including elevated cytochrome P450 expression (17).

To preserve these resistance traits, insecticide-resistant strains were periodically exposed to 0.05% deltamethrin-treated papers every 3–5 generations, following the WHO susceptibility bioassay protocol. Annual bioassays were conducted to assess susceptibility to a range of insecticides, including pyrethroids, carbamates, and organophosphates. Molecular genotyping was also routinely performed to verify the presence of kdr (L1014F) mutations and elevated cytochrome P450 expression, ensuring the stability of resistance mechanisms over time. Further details on strain origins and maintenance protocols are available in (17).

Experiments, referred to as trials, were conducted between June 2019 and February 2020 in a custom-built, climate-controlled free-flight testing room (7 × 4.8 m, 2.5 m high). Trials took place in the afternoon to align with the ‘night’ phase of the mosquitoes’ circadian rhythm, corresponding to their natural host-seeking behaviour. Mosquitoes were tracked using paired recording systems, each comprising a 12 MPixel Ximea CB120RG-CM camera with a 14 mm focal length lens. Each camera was aligned with a Fresnel lens (1400 × 1050 mm, 3 mm thick, 1.2 m focal length; NTKJ Co., Ltd, Japan) positioned approximately 1210 mm away. Retroreflective screens (2.88 m^2^, coated with high-gain sheeting) were placed behind the bednet to enhance light capture and improve contrast for video tracking (18). Infrared illumination (850 nm wavelength) enabled night-phase tracking without disturbing the mosquitoes, capturing trajectories at 50 frames per second.

The system produced 2D telecentric data on mosquito flight patterns. Further details on the experimental setup can be found in (10), and trajectory extraction methods are described in (18). Table 1 summarises the dataset, including a minimum track duration threshold of 1 second.

**Table 1:** Information on the trajectories of each mosquito strain.

| Strain | Bednet Type | Insecticide-resistance Status | Number of Tracks | Average Track Duration (s) | Average Velocity (m/s) |
| --- | --- | --- | --- | --- | --- |
| Kisumu | Untreated | Susceptible | 7,631 | 22.39 (1.00-698.11) | 344.29 (5.20-2,290.84) |
| N’goussu | Untreated | Susceptible | 6,125 | 21.40 (1.00-1,010.03) | 453.78 (3.54-2,602.18) |
| Banfora | Untreated | Resistant | 7,189 | 24.26 (1.00-475.42) | 389.57 (0.74-2,478.15) |
| VK7 | Untreated | Resistant | 5,862 | 17.04 (1.00-457.77) | 429.54 (4.10-2,102.89) |
| Kisumu | Olyset | Susceptible | 936 | 18.59 (1.00-317.98) | 504.43 (4.33-2,709.62) |
| N’goussu | Olyset | Susceptible | 1,801 | 8.77 (1.00-227.30) | 658.58 (5.89-2,819.28) |
| Banfora | Olyset | Resistant | 2,088 | 10.59 (1.00-336.33) | 589.27 (3.78-2,977.75) |
| VK7 | Olyset | Resistant | 1,986 | 10.14 (1.00-341.11) | 554.50 (1.74-3,273.68) |

### 2.2 Data Processing

The dataset contains trajectories of varying durations, which can bias ML model by impacting the features generated. This variation in duration can result due to small obstructions in the camera view, the mosquitoes walking or resting on the bednet or mosquitoes leaving the camera view, leading to the trajectories to be broken.

To account for this, we utilised a moving window technique to split tracks into segments and unify track durations (13). The window size and window overlap of this windowing technique were then optimised for the pipeline through hyperparameter tuning described in section 2.9.

Due to positions being missed, linear interpolation was applied to maintain track continuity. However, in some cases where there were many consecutive interpolated positions or large gaps in tracks, the information quality was reduced.

To address this issue, a segment quality metric was used to score each trajectory segment, where a larger score denotes a lower information quality segment (14). A segment quality threshold was defined to remove segments that are higher than this. To define the threshold, the mutual information of each segment at various quality levels was computed. The weighted average of the thresholds for each segment feature that maximised the information content was used as the final threshold. Segments with quality scores larger than this threshold were removed, resulting in a filtered dataset.

### 2.3 Feature Extraction

Features of flight were extracted from the trajectories. Some features describe the kinematic nature of the tracks, capturing the movement dynamics such as the speed, acceleration and turning angles of the mosquito. The other features describe the geometric characteristics and shape of the track, such as the curvature of the track. A statistical summary was computed for each feature for each trajectory segment. The statistical summary only considers real positions (i.e., not interpolated) within a segment. The calculations for the features of flight are detailed in (13), (14) and provided in the supplementary.

### 2.4 Dataset Partitioning

The dataset, containing 40 total experiment trials (23 Olyset trials and 17 Untreated trials), was randomly split into two distinct datasets: a modelling set and a tuning set. The tuning set was used for feature selection and hyperparameter tuning, consisting of 19 trials. The modelling set, containing 21 trials, was dedicated to the ML model training and evaluation. Partitioning the dataset by trials prevents data leakage and better reflects real-world scenarios where entire trials are tested independently. This approach enables robust model training, ensures thorough cross-validation, and maintains the reliability of performance evaluations.

### 2.5 Feature Selection

To select meaningful features, the Mann-Whitney U-test, a non-parametric statistical test, from SciPy (19) was used. A family-wise error rate (FWER) controlling procedure using Bonferroni correction was used to reduce group testing p-value inflation. Features that rejected the null hypothesis at an FWER *<* 0.05 were selected. Highly correlated features were removed by calculating the Spearmans correlation and removing one feature from each pair of highly correlated features (*ρ >* 0.85) from the set of selected features.

### 2.6 Classification Model

This study aims to differentiate between behaviours of mosquitoes in the presence of an untreated net and the Olyset ITN using a ML classifier. The XGBoost classifier (20), a boosting-tree algorithm, known for being the state-of-the-art in tabular classification, was used. In order to generate whole track predictions, every track segment was classified independently and the mode of the segment binary predictions for each track was subsequently used as the prediction for the complete track.

Prior to training, each feature was standardised using Z-score normalisation, where the mean and standard deviation values were calculated from the training set. As the training dataset may have class imbalance, the scale pos weight parameter within the XGBoost model was adjusted to be the ratio of the number of UT segments to the number of OL segments.

The decision boundary of the model was also adjusted. Typically, the predictions of a model are considered through the probability score of each class, where in a binary classification task, a probability score above 0.5 for a given class leads to that class being the final prediction. The decision boundary is adjusted by evaluating the training dataset at varying decision boundaries { 0.01, 0.02, …, 0.99}. The decision boundary that returns the largest Matthews correlation coefficient score on the training set is used to set the decision boundary threshold on the test.

### 2.7 Model Training Strategies

The ML model was trained using a subset of trials. Each trial consists of a 2-hour long experiment where mosquito trajectories were generated. For insecticide-susceptible strains on the Olyset net, it was noticed that mosquitoes were killed or highly impacted after 30 minutes. As such, there were very few tracks generated after 30 minutes in the experiment, and those that exist were highly distorted and unreliable.

To ensure the ML model was trained on reliable and consistent trajectories, a balanced training strategy (BTS) was employed. This involved training the model using OL IS tracks generated within 30 minutes of the experiment as well as sampling OL IR, UT IS and UT IR tracks so that the numbers of tracks for each class are approximately equal. Other training strategies were also considered such as an early training strategy (training the model using tracks generated within 30 minutes of each experiment) and comprehensive training strategy (training the model on all available data). Results of these other training strategies can be found in the supplementary and were similar or slightly worse than the BTS.

### 2.8 Evaluation and Interpretation

The ML model was evaluated on the modelling dataset using a cross-validation approach. Within cross validation, the ML model is iteratively trained and tested on a different subset of training and testing datasets. Each iteration is known as a fold, with the training set consisting of 1 UT trial from each strain, and 2 OL trials from each strain (see section 2.1). This was done so that the number of tracks in each class was more balanced, as the number of trajectories within the OL class is much less per trial (see Table 1). The other trials are used in training. Overall, there are 1944 possible folds, with which 30 combinations were sampled. The training set thus usually consists of 12 training trials and 9 testing trials.

Performance metrics were calculated on whole tracks using Scikit-learn (21), including balanced accuracy, ROC AUC (area under the receiver operator curve) score, PR AUC (area under the precision-recall curve) score, F1 score, precision score, recall score, Matthew Correlation Coefficient (MCC), Cohen kappa coefficient and log-loss score. The arithmetic mean, minimum and maximum of the performances across all folds was provided.

Confusion matrices provided a visual summary of model performance, showing the normalised percentages of true and false predictions for IR and IS classifications relative to the true labels. High diagonal values indicated correct classifications, while off-diagonal values highlighted errors. Under random chance, each row would display a 50-50 split between classes. Normalising the data made it easier to evaluate performance within each class and identify misclassification patterns.

To explain the model predictions, Shapley Additive ExPlanations (SHAP) (15) values are used. These represent the contribution of each feature to predictions made by the ML model. By using this, we can gain insights into the features that have the strongest influence on the model’s output and how they interact with each other. Here, the SHAP values are calculated on track segments, which can aid the understanding of the differences between mosquito behaviours around untreated bednets and the Olyset net.

The impact of insecticide on IS strains acts fairly quickly, thus it is necessary to explore the changes in accuracy of the ML model across the experiment time. Each trial is 2 hours long, so the accuracy for every 5-minute segment of the experiment can be plotted. For tracks that starts within the 5-minute segment, the average accuracy was computed and plotted as a line graph. Similarly, performance metrics are calculated for each strain, each class, and before/after 30 minutes of the experiment.

### 2.9 Hyperparameter Tuning

The ML pipeline has various parameters that require optimising to obtain the best performance whilst also reducing overfitting. This includes the window size and overlap of the windowing technique used to split tracks into segments, as well as the ML model parameters. The parameters were tuned together in a cross-validated grid search approach attempting to maximise balanced accuracy on an unseen test set. A full description of the parameter ranges and step sizes, as well as the full set of optimised hyperparameters identified for each model can be found in the supplementary information.

## 3 Results

### 3.1 Data Processing

Through hyperparameter tuning, a segment size of 7.5 seconds and a segment overlap of 7 seconds obtained the best performance on the tuning data set. Using these parameters, the final dataset contains 12,991 tracks, which are split into 881,713 track segments.

### 3.2 Model Evaluation

Table 2 provides model performance metrics, including the arithmetic mean, minimum, and maximum values for each metric (shown in brackets). Performance indicators include F1 score, recall, precision, and the area under the precision-recall curve (PR AUC), with calculations made for both UT and OL as the positive class. These summary statistics offer a clear view of the model’s variability and stability across different evaluation metrics.

**Table 2:** Performance metrics of the XGBoost classification model on independent test data. Each evaluation metric is reported as the arithmetic mean, with the minimum and maximum values provided in brackets across all crossvalidation folds.

| Evaluation Metric | Performance |
| --- | --- |
| Balanced accuracy | 0.838 (0.798 - 0.888) |
| ROC AUC | 0.925 (0.899 - 0.944) |
| Matthew Correlation Coefficient | 0.502 (0.442 - 0.574) |
| Log loss | 0.354 (0.288 - 0.460) |
| Cohen Kappa Coefficient | 0.465 (0.371 - 0.558) |
| F1 score (OL) | 0.521 (0.427 - 0.606) |
| F1 score (UT) | 0.935 (0.900 - 0.953) |
| Recall (OL) | 0.782 (0.683 - 0.908) |
| Recall (UT) | 0.894 (0.824 - 0.939) |
| Precision (OL) | 0.397 (0.283 - 0.538) |
| Precision (UT) | 0.979 (0.967 - 0.991) |
| PR AUC (OL) | 0.489 (0.395 - 0.621) |
| PR AUC (UT) | 0.803 (0.771 - 0.839) |

Figure 1 contains the ROC curves with each folds’ ROC curve shown as a different coloured line, as well as the confusion matrix for the balanced method. As the test set was imbalanced, the confusion matrix shows the normalised percentages of the predictions across all folds. The normalisation was performed over the true class such that the proportion of correct predictions can be highlighted.

**Fig. 1:**
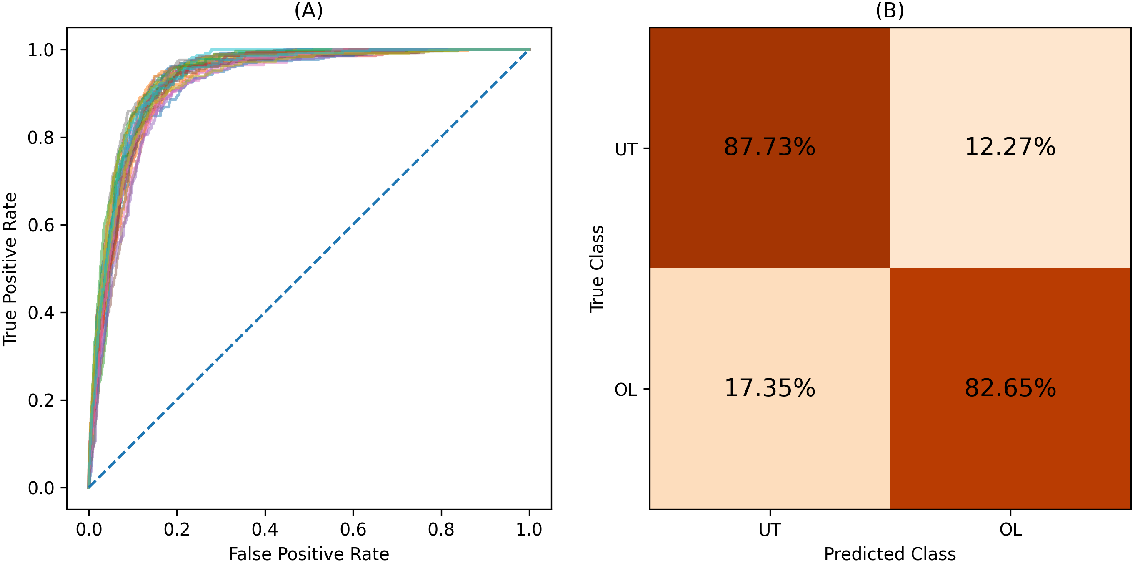
Model performance evaluation. (A) Receiver Operating Characteristic (ROC) curves for each fold, illustrating the trade-off between sensitivity (true positive rate) and 1-specificity (false positive rate) across classification thresholds. Each line represents the performance of an individual fold, providing insight into model consistency. (B) Confusion matrix derived from independent test data, displaying the normalised percentages of predictions averaged across all folds. The matrix is normalised by the true class, highlighting the proportion of correct and incorrect predictions within each class.

To examine how classifier performance changes over time, we plotted the average accuracy within each 5-minute segment of the 2-hour experimental trials for both the OL and UT nets. Figure 2 shows accuracy trends for the OL net, while Figure 3 shows trends for the UT net. In these plots, the central line represents the average accuracy across all folds for track segments that start within each 5-minute interval. The surrounding blue shaded area represents the standard deviation, highlighting the variation in accuracy across folds.

**Fig. 2:**
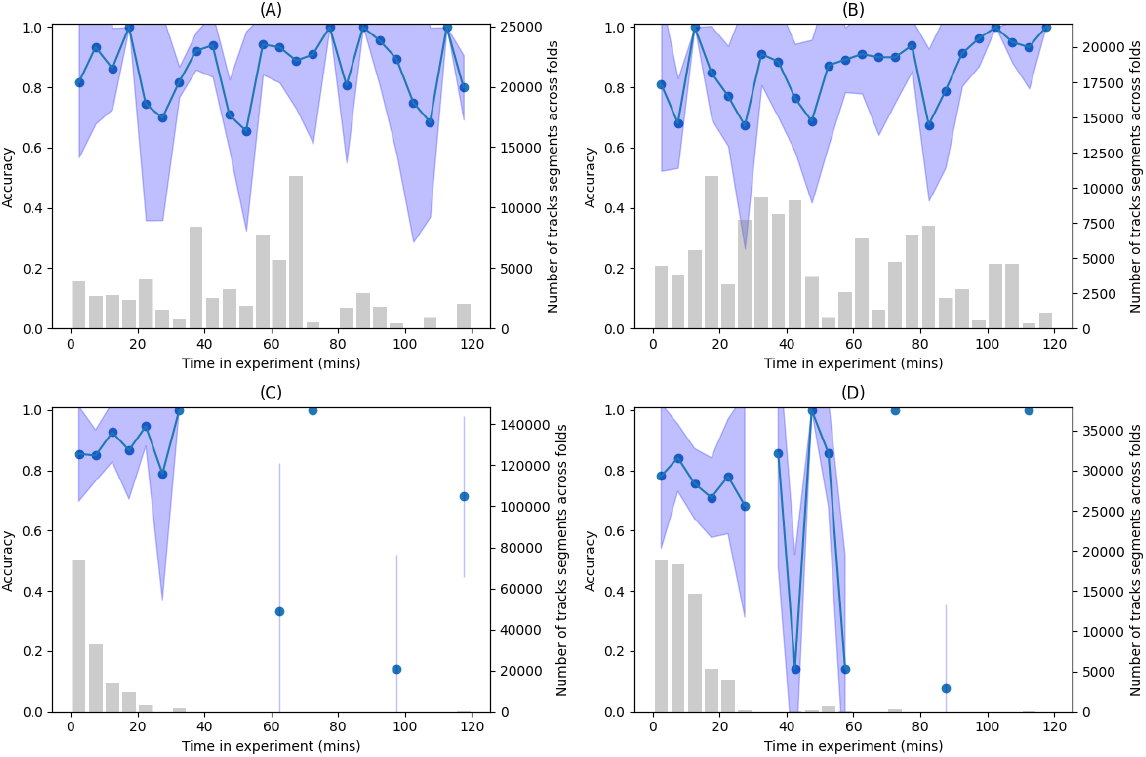
Model performance across experiments for each mosquito strain on the Olyset net: (A) Banfora, (B) VK7, (C) Kisumu, and (D) N’goussu. The solid lines and dots represent the average classification accuracy of the model, while the shaded blue regions indicate the standard deviation across all cross-validation folds, providing a measure of variability and robustness. The accompanying bar plots illustrate the total number of track segments analyzed across all folds for each strain, highlighting the dataset size and distribution.

**Fig. 3:**
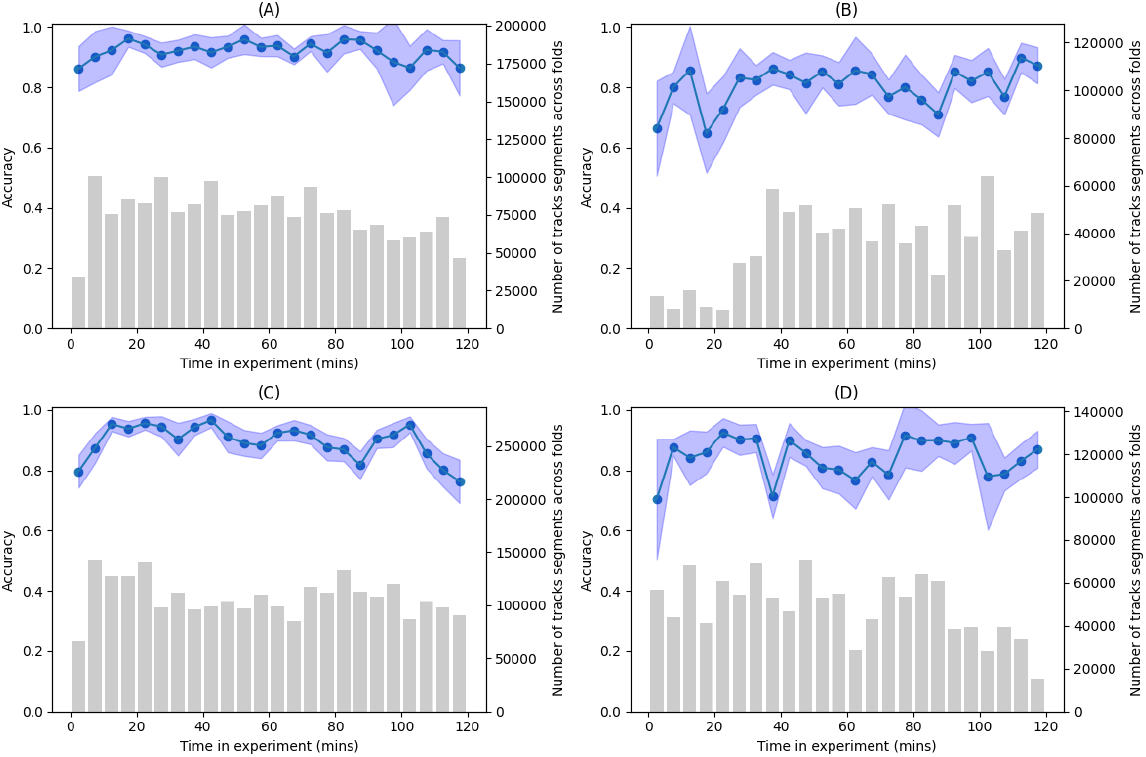
Model performance across experiments for each mosquito strain on the untreated net: (A) Banfora, (B) VK7, (C) Kisumu, and (D) N’goussu. The solid lines and dots represent the average classification accuracy of the model, while the shaded blue regions indicate the standard deviation across all cross-validation folds, providing a measure of variability and robustness. The accompanying bar plots illustrate the total number of track segments analysed across all folds for each strain, highlighting the dataset size and distribution.

For the best performing fold, SHAP plots are provided in Figure 4. This includes a SHAP bar plot, and SHAP summary plot. SHAP summary and bar plots for the other training strategies are provided in the supplementary and display the same trends.

**Fig. 4:**
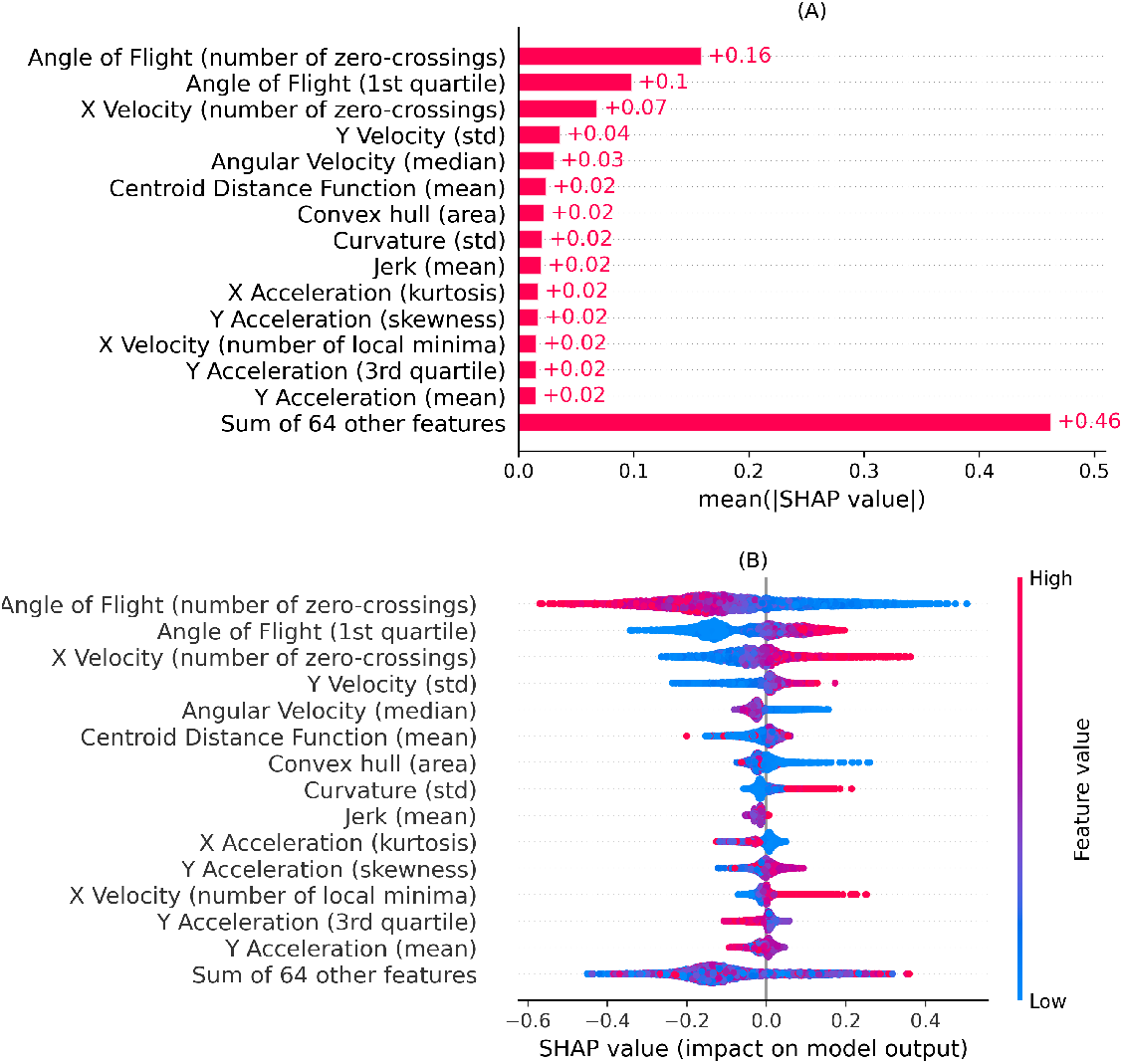
SHAP visualisations for the best-performing model fold applied to independent test data. Panel (A) shows the SHAP bar plot, which ranks features based on their mean absolute SHAP values, indicating their overall importance in the model’s predictions. Panel (B) presents the SHAP summary plot, where each dot represents an individual data segment. The colour gradient of the dots reflects the feature values, with red indicating higher values and blue indicating lower values. In both visualizations, features are sorted by their mean absolute SHAP values. Positive SHAP values indicate contributions towards the OL class, whereas negative SHAP values indicate contributions towards the UT class. These plots provide insights into feature influence and directionality within the classification model.

## 4 Discussion

ITNs play a pivotal role in controlling the spread of mosquito borne diseases like malaria. However, understanding mosquito behaviour in response to insecticides and how these behaviours differ between resistant and susceptible strains can reveal potential areas for improvement in vector control strategies. Our machine learning models displayed an outstanding ability to distinguish mosquito trajectories captured around an untreated net and a treated net, providing insight into the behavioural responses and adaptions made by mosquitoes when around an ITN. Across the 3 training approaches, model performance remained exceptional with average balanced accuracies between 0.813 and 0.838, and average ROC AUC scores exceeding 0.92. The un-normalised confusion matrices (see Supplementary) appear distorted due to imbalance in the test datasets, as a result of IS mosquitoes at the OL net being killed after 30 mins whereas IR mosquitoes are present throughout. It follows that over the 2-hour experiments, there is little-to-no variation for UT accuracy, whereas OL accuracy displays more varied performance, shown in Figures 2 and 3. It was hypothesised that the accuracy on the OL net would be poor initially and then improve with time as the behaviour of the mosquitoes would change from UT to OL. However, the results demonstrate that there is an immediate behavioural response at an OL net for both IR and IS mosquitoes. This could suggest that net contact occurs very quickly as the mosquitoes are attracted to the human bait, or that the OL net has repellent properties (discussed below).

The SHAP analysis identified three dominant features contributing to model predictions: the number of zero-crossings in flight angle change, the 1st quartile of flight angle change, and the number of zero-crossings in horizontal velocity. Together, these features suggest that mosquitoes interacting with the Olyset net experience highly convoluted flight paths, and are likely indicative of disorientation caused by insecticidal exposure.

As the change in angle of flight is always a positive value, the number of zero-crossings in this feature represents the total number of zero values in a track segment. In other words, a high value represents little change in angle within a segment. In the SHAP summary plot (Figure 4), the trajectory segments with low numbers of zero crossings in angle of flight change tend to contribute towards the OL class – suggesting that tracks on the OL net are more convoluted than UT. Similarly, the higher values of 1st quartile of angle of flight change, indicates that larger angle changes were experienced within a track segment, and tended towards the OL class. Different to change in angle of flight, horizontal velocity is a signed quantity, and thus the number of zero crossings represents the total number of zero values and zero-crossings. A zero crossing in this case means that the mosquito has changed direction in the X direction (across the camera). Larger values of this feature also tend towards the OL net class, where high values suggest more direction changes. These three features account for most of the predictive power of the ML model. Across them all, they suggest that track segments on the OL net are highly convoluted with many large angle changes, as well as more direction changes. Simply put, these tracks are more chaotic as the mosquitoes are impacted by the insecticide.

Previous studies demonstrate conflicting evidence of the repellency of permethrin the active ingredient in the Olyset net. For example, it was found that permethrin-treated clothing repelled 40-52% of Aedes albopictus mosquitoes (22) whereas in an experimental hut trial the results for Anopheles gambiae at Olyset nets showed the mosquitoes were not repelled (23). Some of the confusion has arisen from the use of differing definitions for the behavioural reactions of mosquitoes to insecticide (24). Here, we define repellency as a pre-contact response, where the presence of the insecticide deters mosquitoes from approaching the treated net. Irritancy, on the other hand, we define as a behavioural response after contact with insecticide-treated surfaces, which may potentially cause disorientation in the mosquito. In the literature permethrin was found to be repellent in cases where a permethrin solution was created to coat or impregnate a material, either a bednet (25), test papers (22) or textile samples (26). In particular, Lindsay et al (27) considered the observed repellency to be associated with volatile constituents of the emulsifier concentrate rather than permethrin itself. In contrast, when the permethrin was fused with resin to form the fibres used in the Olyset ITN, a number of studies failed to find evidence for repellency, e.g. Spitzen et al (23) and Mueller et al (24). Our model reveals that mosquitoes display sustained disrupted flight behaviour at the OL net compared to a UT net. For individual segments the classifier exhibits higher noise and it has not been possible to reliably examine changes in behaviour before and after a mosquito’s first contact with the Olyset ITN. Therefore, the effective temporal resolution of this ML approach is the trajectory length which has average values between 10.14 and 24.26 seconds, see Table 1. This period may include first contact with the net and hence a small contribution of repellent behaviours as well as irritancy. Hence, further studies need to be conducted to identify any repellent properties.

Both resistant and susceptible mosquitoes exhibit similarly disrupted flight behaviours around the Olyset net. This is a key finding, as it suggests that resistance may not entirely shield mosquitoes from the behavioural effects of the insecticide, even if it improves their survival. Both IR and IS mosquitoes exhibit convoluted and erratic flight trajectories when exposed to the Olyset net, characterised by frequent direction changes and chaotic movements (see Figure 10 in supplementary).

Interestingly, within the first 30-minutes of the experiment the number of IS trajectory segments at the Olyset net outnumbered those of IR (even though the same number of mosquitoes are released for each trial). The filtered and unfiltered segments exhibit the same trend. The duration and distance travelled by the IS mosquitoes at the OL net are less than those for IR and are shorter than those for IS mosquitoes at an UT net (see boxplots in supplementary). This suggests that IS mosquitoes are disproportionately affected by the insecticide in the early stages, exhibiting more rapid flight paths, possibly in attempts to escape or avoid the net before dying due to insecticide toxicity.

Three distinct training strategies were employed to establish models for mosquito behaviour at the OL and UT nets. The comprehensive strategy included late-stage behaviour when insecticidal effects were at their peak, while the early training strategy excluded these to focus on periods where both IR and IS mosquitoes are active. The balanced training strategy (BTS) attempts to equalise the number of tracks between IR and IS mosquitoes. The focus has been given to the results generated via the BTS which yielded the best performance, nevertheless, the other training strategies returned similar conclusions.

While insecticide resistance continues to challenge the efficacy of traditional vector control tools (4), our findings indicate that exposure to single-ingredient pyrethroids can alter mosquito flight patterns, even in resistant populations. These observations highlight the potential for leveraging behavioural responses in the design of novel vector control strategies. The ML approach demonstrated here provides a scalable framework for evaluating compounds based on their effects on mosquito behaviour, enabling efficient screening prior to field testing.

While our ML approach revealed novel insights into mosquito behavioural responses to Olyset nets, several limitations warrant consideration in future research. The study focused specifically on Anopheles gambiae and permethrin-treated nets - future work should be expanded to analyse other important vector species (e.g., the more exophilic sibling species An arabiensis) or diurnally active mosquitoes, like the arbovirus vector Aedes aegypti, and alternative insecticide classes (or combinations thereof) and the surfaces or materials to which they are applied (e.g. the solid surfaces used for indoor residual praying) to develop a comprehensive understanding of multiple behavioural resistance mechanisms. Additionally, while our ML model successfully classified flight patterns, underlying physiological mechanisms driving these behavioural changes remain unclear. Further research is needed to investigate these mechanisms and the factors driving behavioural responses to insecticides.

