## Supplementary Information for "Machine Learning Reveals Immediate Disruption in Mosquito Flight when exposed to Olyset Nets"

Y.M. Qureshi[1], V. Voloshin[1, 2], A. Guy[3], H. Ranson[3], P.J. McCall[3],

J.A. Covington[1], C.E. Towers[1], D.P. Towers[1]

[1]School of Engineering, University of Warwick, Coventry, CV4 7AL, UK

[2]School of Biological and Behavioural Sciences, Queen Mary University of London, E1 4NS, UK

[3]Vector Biology Department, Liverpool School of Tropical Medicine, Pembroke Place, Liverpool, L3 5QA, UK

Corresponding Author:

 (YMQ)

### Feature Calculations

1. Velocity, acceleration and Jerk

These are calculated using second-order central finite difference methods. The signed values are used – where in the case of velocity, a positive velocity indicates motion from head-to-toe of the human bait within the experiment. The axial values are also utilised.

2. Angle of Flight

The angle of flight, $\alpha_i$, can be calculated using the dot product of two vectors. It describes the relative change in direction between positions.

$$v_i = \begin{bmatrix} x_i - x_{i-1} \\ y_i - y_{i-1} \end{bmatrix}$$

$$v_{i+1} = \begin{bmatrix} x_{i+1} - x_i \\ y_{i+1} - y_i \end{bmatrix}$$

$$\alpha_i = \arccos\left(\frac{v_i \cdot v_{i+1}}{\sqrt{(v_i \cdot v_i)(v_{i+1} \cdot v_{i+1})}}\right)$$

3. Angular Velocity and Angular Acceleration

The angular velocity and angular acceleration are both calculated using second-order finite difference methods. The arctangent function is used to calculate the angle between positions, then the change in angle between consecutive positions is computed. Angular velocity is calculated by dividing this change in angle by the corresponding time difference.

4. Orthogonal Components of Velocity

This is a group of features that describe the tendency of a movement to head in a given direction. The persistence velocity describes the tendency to head tangential to the trajectory while the turning velocity describes the tendency to head normal to the trajectory. It involves converting from the Cartesian to the circular coordinate system:

$$\rho_i = \sqrt{(x_{i+1} - x_i)^2 + (y_{i+1} - y_i)^2}$$

$$\theta_i = \arctan\left(\frac{y_{i+1} - y_i}{x_{i+1} - x_i}\right)$$

The change in $\theta$ and instantaneous velocity is calculated:

$$\Theta_i = |\theta_{i+1} - \theta_i|$$

$$v_i = \frac{\rho_i}{t_{i+1} - t_i}$$

Then converting back to the Cartesian coordinate system:

$$P_i = v_i \cos \Theta$$

$$T_i = v_i \sin \Theta$$

Where $P_i$ and $T_i$ are the persistence and turning velocities, respectively.

5. Tortuosity

This is the ratio of the actual distance travelled and the shortest distance between the start and end positions.

$$S = \frac{\sum_{i=0}^{N} \sqrt{(x_{i+1} - x_i)^2 + (y_{i+1} - y_i)^2}}{\sqrt{(x_N - x_0)^2 + (y_N - y_0)^2}}$$

6. Convex Hull

The convex hull is the set of points that forms the smallest possible convex polygon that bounds all the points of a track. From the polygon that is formed, the area and perimeter is extracted and used as features.

7. Centroid Distance Function

This calculated the distances of each position in a track to the centre point of the track.

$$x_c = \frac{1}{N} \sum_{i=0}^{N} x_i$$

$$y_c = \frac{1}{N} \sum_{i=0}^{N} y_i$$

$$C_i = \sqrt{(x_i - x_c)^2 + (y_i - y_c)^2}$$

8. Curvature

Curvature is a measure that defines the deviation of a trajectory from a straight line. It is calculated as:

$$k_i = \frac{\dot{x}_i \ddot{y}_i - \dot{y}_i \ddot{x}_i}{\left(\dot{x}_i^{\,2} + \dot{y}_i^{\,2}\right)^{\frac{3}{2}}}$$

9. Curvature Scale Space

This measures the inflection points in a trajectory at difference scales and is robust to affine transformations. To compute the curvature at varying levels, the trajectory is convolved with a 1D gaussian kernel of width $\sigma$. The value of $\sigma$ is varies, and the zero-crossings locations in the curvature are recorded. The locations of zero-crossings at different $\sigma$ levels can be plotted as a CSS image. A 1D signal is generated by taking column maximums from this CSS image.

10. Fractal Dimension

Fractal dimension is a measure of the complexity of a trajectory. Values around 1 describe a linear path with a shape similar to a straight line, and values around 2 indicate convoluted movement with a shape similar to a plane. To compute fractal dimension, $d$, the following equations are used:

$$d = \frac{\log n}{\log \frac{1}{S}}$$

Where $n$ is the number of minature pieces in the path, $S$ is the scaling factor, and $d$ is the fractal dimension. For a trajectory, the values of $n$ and $S$ are defined as follows:

$$n = \sum_{i=0}^{N} \sqrt{(x_{i+1} - x_i)^2 + (y_{i+1} - y_i)^2}$$

$$S = \frac{1}{\sqrt{(\max x - \min x)^2 + (\max y - \min y)^2}}$$

**Penalty functions and thresholds assessed.**

To remove tracks with substantial gaps (many missing positions), a segment quality metric was used to score track segments. A mutual information-based method was used to obtain a threshold for the segment quality metric.

This is where we developed a penalty function to score a track on its information content. Let $segment$ be a list of length $N$ where each element in the list is either 1 or 0, indicating a real position (1) or an interpolated position (0) for a segment. The penalty function $P(segment, n, m)$ is defined as:

$$P(segment, n, m) = \frac{1}{N}\sum_{i=0}^{N}\begin{cases} n \cdot m^{c_i}, & \text{if } x_i = 0 \\ 0, & \text{if } x_i = 1 \end{cases}$$

Where $c_i$ is the length of the consecutive artificial positions starting from position $i$ until a real position is encountered. If $x_i = 1$, $c_i$ is reset to 0. Various values of $n$ and $m$ were assessed with $n = 1$ and $m = 1.05$ being used for the final model.

To obtain a threshold for the penalty score, a threshold was determined by computing the mutual information at various score thresholds. Then the weighted average threshold is obtained based on the maximum mutual information and its corresponding threshold for each feature. This weighted average is the final score threshold.

Table 1. Hypertuning parameter ranges

| Parameter | Parameter values |
|---|---|
| Window size | 0.5, 1, 1.5, 2, 2.5, 3, 3.5, 4, 4.5, 5, 5.5, 6, 6.5, 7, 7.5, 8, 8.5, 9, 9.5 |
| Window overlap | 0.5, 1, 1.5, 2, 2.5, 3, 3.5, 4, 4.5, 5, 5.5, 6, 6.5, 7, 7.5, 8, 8.5, 9 |
| Learning rate | 0.01, 0.1, 0.3 |
| N estimators | 50, 100, 150, 200, 250 |
| Max depth | 3, 5, 7 |
| Subsample | 0.5, 0.7, 0.9 |
| Colsample bytree | 0.5, 0.7, 0.9 |
| Reg alpha | 0, 0.01, 0.1 |
| Reg lambda | 0, 0.01, 0.1 |
| Min child weight | 1, 5, 15 |

Table 2. Final hyperparameters for each training strategy.

| Parameter | Comprehensive | Balanced | Early |
|---|---|---|---|
| Window size | 7 | 7.5 | 8 |
| Window overlap | 6.5 | 7 | 7 |
| Learning rate | 0.3 | 0.3 | 0.3 |
| N estimators | 200 | 100 | 200 |
| Max depth | 3 | 5 | 3 |
| Subsample | 0.8 | 0.7 | 0.8 |
| Colsample bytree | 0.9 | 0.8 | 0.9 |
| Reg alpha | 0 | 0.1 | 0.1 |
| Reg lambda | 0 | 0.1 | 0 |

| | | | |
|---|---|---|---|
| Min child weight | 10 | 20 | 10 |

Table 3. Performance metrics for the comprehensive and early data training strategies.

| Performance metric | Comprehensive data training | Early data training |
|---|---|---|
| Balanced accuracy | 0.813 (0.769 - 0.863) | 0.838 (0.785 - 0.885) |
| ROC AUC | 0.931 (0.903 - 0.952) | 0.925 (0.897 - 0.945) |
| Matthew Correlation Coefficient | 0.521 (0.458 - 0.618) | 0.498 (0.432 - 0.562) |
| Log loss | 0.323 (0.287 - 0.361) | 0.360 (0.290 - 0.453) |
| Cohen Kappa Coefficient | 0.506 (0.441 - 0.615) | 0.459 (0.356 - 0.556) |
| F1 score (OL) | 0.554 (0.490 - 0.654) | 0.516 (0.414 - 0.603) |
| F1 score (UT) | 0.950 (0.938 - 0.960) | 0.933 (0.902 - 0.953) |
| Recall (OL) | 0.696 (0.595 - 0.809) | 0.785 (0.656 - 0.904) |
| Recall (UT) | 0.929 (0.897 - 0.962) | 0.892 (0.831 - 0.938) |
| Precision (OL) | 0.466 (0.355 - 0.625) | 0.391 (0.273 - 0.534) |
| Precision (UT) | 0.972 (0.958 - 0.987) | 0.979 (0.964 - 0.991) |
| PR AUC (OL) | 0.526 (0.392 - 0.659) | 0.489 (0.392 - 0.612) |
| PR AUC (UT) | 0.799 (0.763 - 0.833) | 0.803 (0.772 - 0.839) |

Table 4. Windowing parameters for each model alongside the final number of tracks and segments.

| Training Strategy | Window Size (s) | Window Overlap (s) | Final number of tracks | Number of segments |
|---|---|---|---|---|
| Comprehensive data training | 7 | 6.5 | 13,469 | 1,067,889 |
| Early data training | 8 | 7 | 12,584 | 440,118 |
| Balanced data training | 7.5 | 7 | 12,991 | 881,713 |

Table 5. Accuracy of the XGBoost classification model when applied to independent test data with differing data training strategies for resistant and susceptible strains, before and after 30 minutes.

| Training strategy | Comprehensive | Early | Balanced |
|---|---|---|---|
| OL IR (<30mins) | 0.921 (0.803 - 1.000) | 0.912 (0.750 - 1.000) | 0.929 (0.812 - 1.000) |
| OL IS (<30mins) | 0.836 (0.647 - 0.958) | 0.844 (0.669 - 0.934) | 0.877 (0.727 - 0.961) |
| OL IR (>30mins) | 0.894 (0.791 - 0.942) | 0.889 (0.805 - 0.967) | 0.905 (0.827 - 0.962) |
| OL IS (>30mins) | 0.705 (0.000 - 1.000) | 0.667 (0.000 - 1.000) | 0.752 (0.000 - 1.000) |
| UT IR (<30mins) | 0.859 (0.816 - 0.904) | 0.866 (0.797 - 0.927) | 0.852 (0.802 - 0.905) |
| UT IS (<30mins) | 0.893 (0.860 - 0.933) | 0.899 (0.865 - 0.936) | 0.883 (0.843 - 0.915) |
| UT IR (>30mins) | 0.854 (0.821 - 0.879) | 0.849 (0.805 - 0.878) | 0.844 (0.800 - 0.877) |
| UT IR (>30mins) | 0.852 (0.814 - 0.873) | 0.834 (0.792 - 0.871) | 0.836 (0.802 - 0.857) |

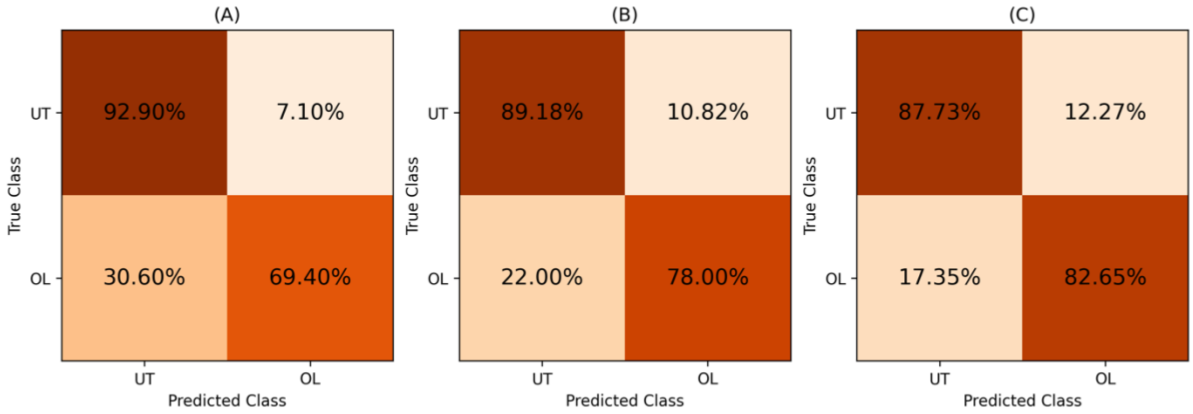


Figure 1. Confusion matrices from independent test data displaying the normalised percentages of the predictions of the true class over all folds. In the figure, (A) displays the comprehensive data training strategy, (B) is the early data training strategy, and (C) is the balanced data training strategy.

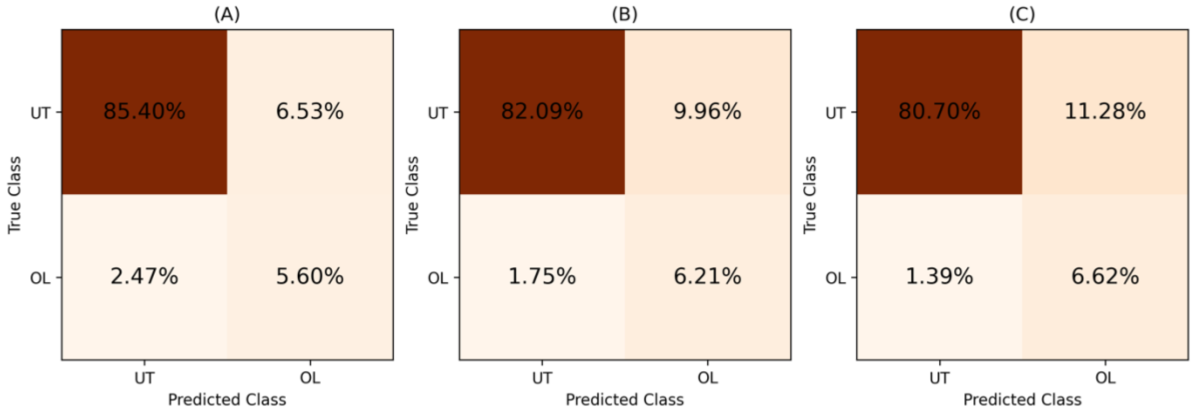


Figure 2. Confusion matrices from independent test data displaying the percentages of the predictions over all folds. In the figure, (A) displays the comprehensive data training strategy, (B) is the early data training strategy, and (C) is the balanced data training strategy.

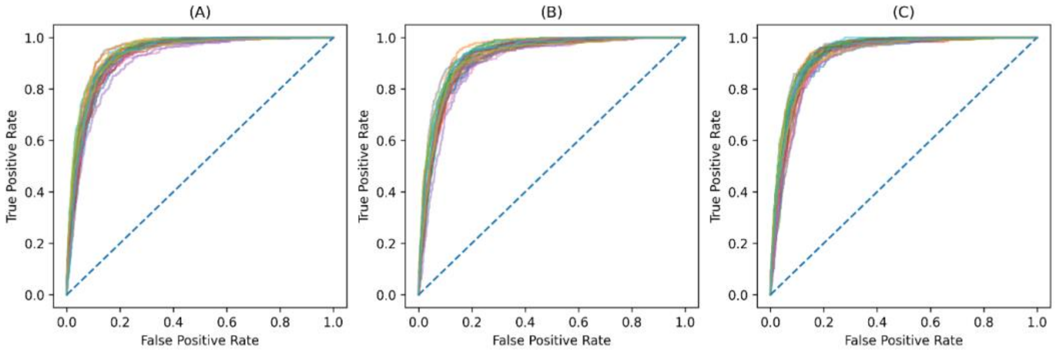


Figure 3. ROC Curves from independent test data displaying different coloured lines for each folds' ROC curve. In the figure, (A) displays the comprehensive data training strategy, (B) is the early data training strategy, and (C) is the balanced data training strategy.

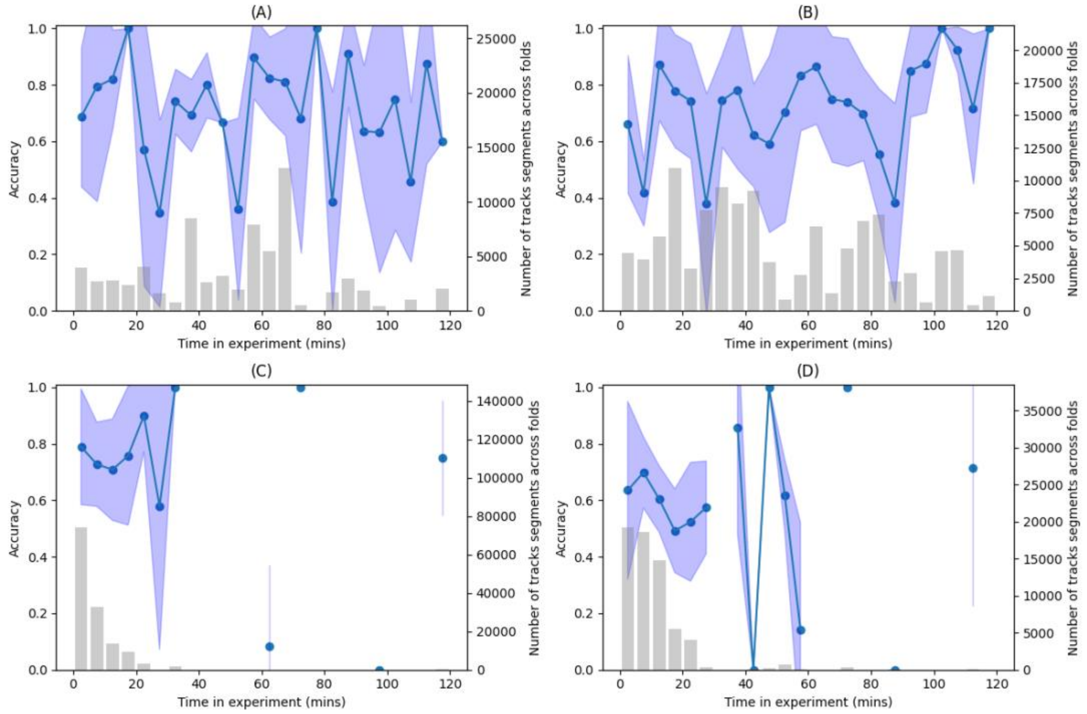


Figure 4. Model performance across the experiments for the comprehensive data training strategy for each strain on the Olyset net – (A) Banfora, (B) VK7, (C) Kisumu, and (D) Ngoussu. The lines and dots indicate the model accuracy in classification, whilst the bar plot displays the sum of the number of tracks across all folds.

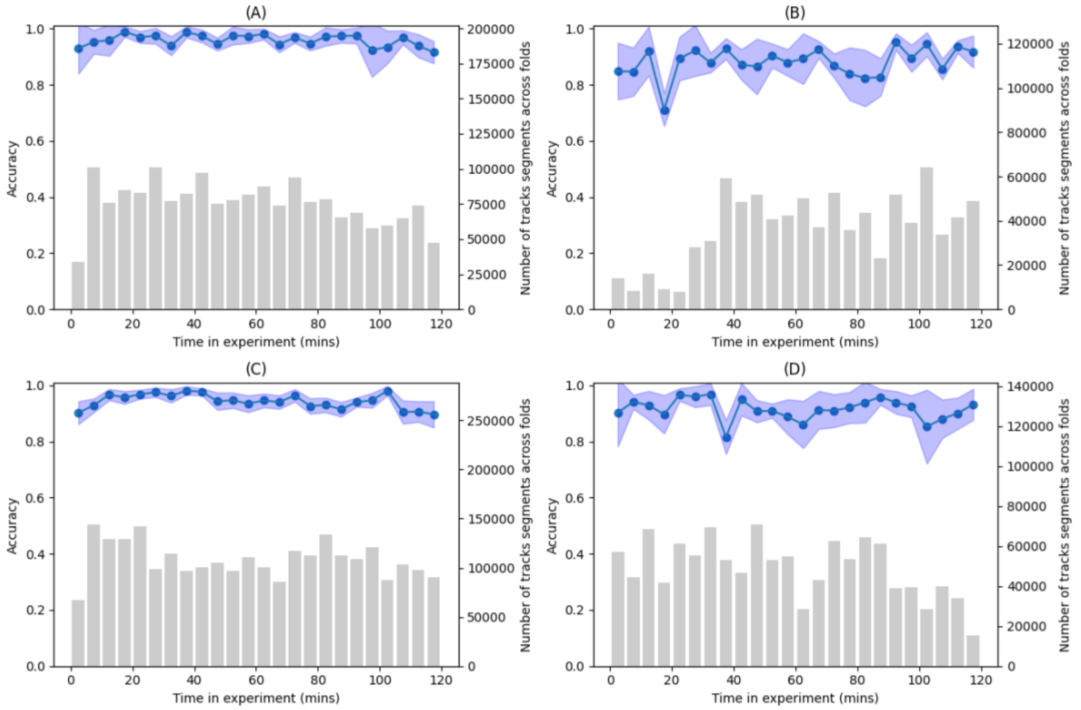


Figure 5. Model performance across the experiments for the comprehensive data training strategy for each strain on the untreated net – (A) Banfora, (B) VK7, (C) Kisumu, and (D) Ngoussu. The lines and dots indicate the model accuracy in classification, whilst the bar plot displays the sum of the number of tracks across all folds.

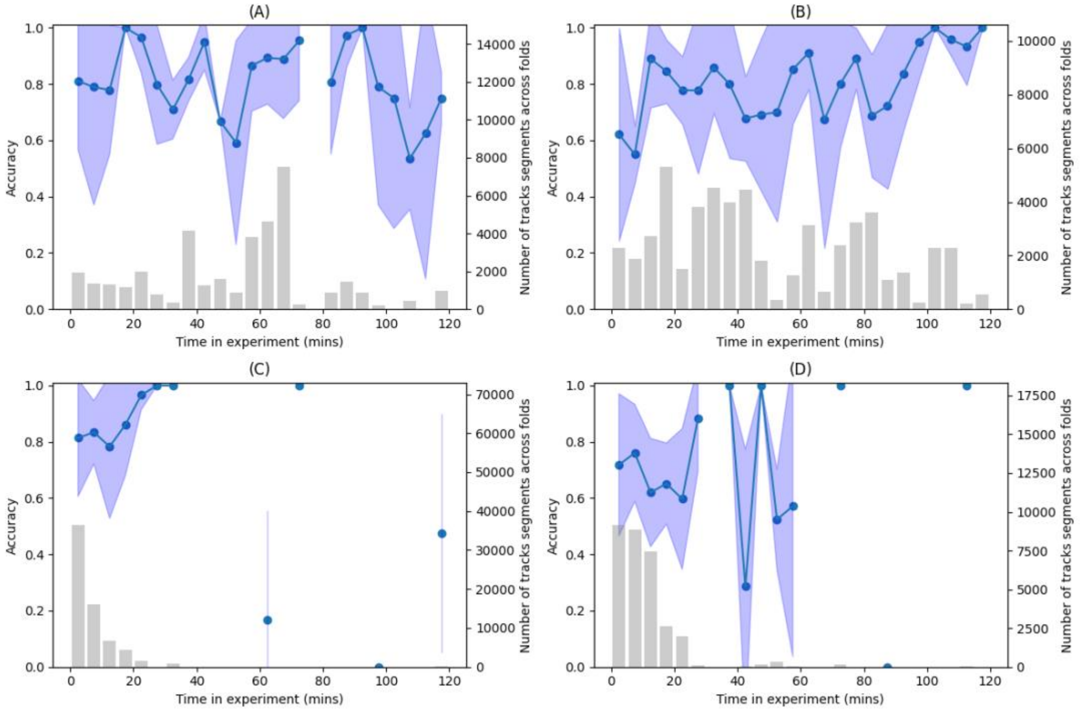

Figure 6. Model performance across the experiments for the early data training strategy for each strain on the Olyset net – (A) Banfora, (B) VK7, (C) Kisumu, and (D) Ngoussu. The lines and dots indicate the model accuracy in classification, whilst the bar plot displays the sum of the number of tracks across all folds.

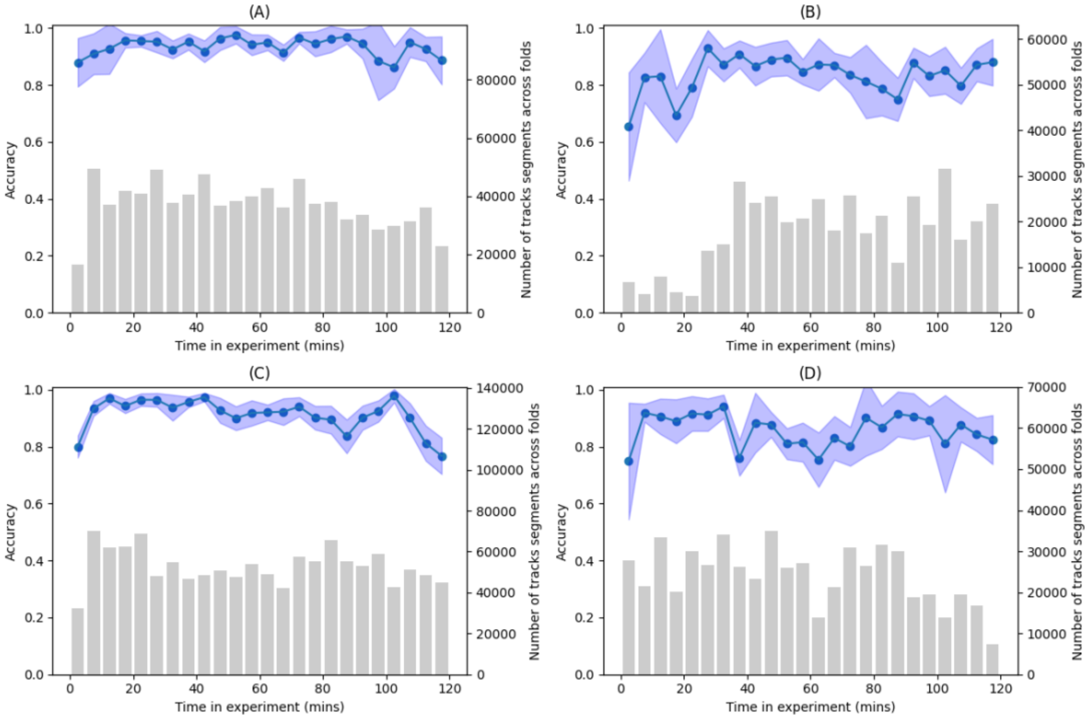


Figure 7. Model performance across the experiments for the early data training strategy for each strain on the Untreated net – (A) Banfora, (B) VK7, (C) Kisumu, and (D) Ngoussu. The lines and dots indicate the model accuracy in classification, whilst the bar plot displays the sum of the number of tracks across all folds.

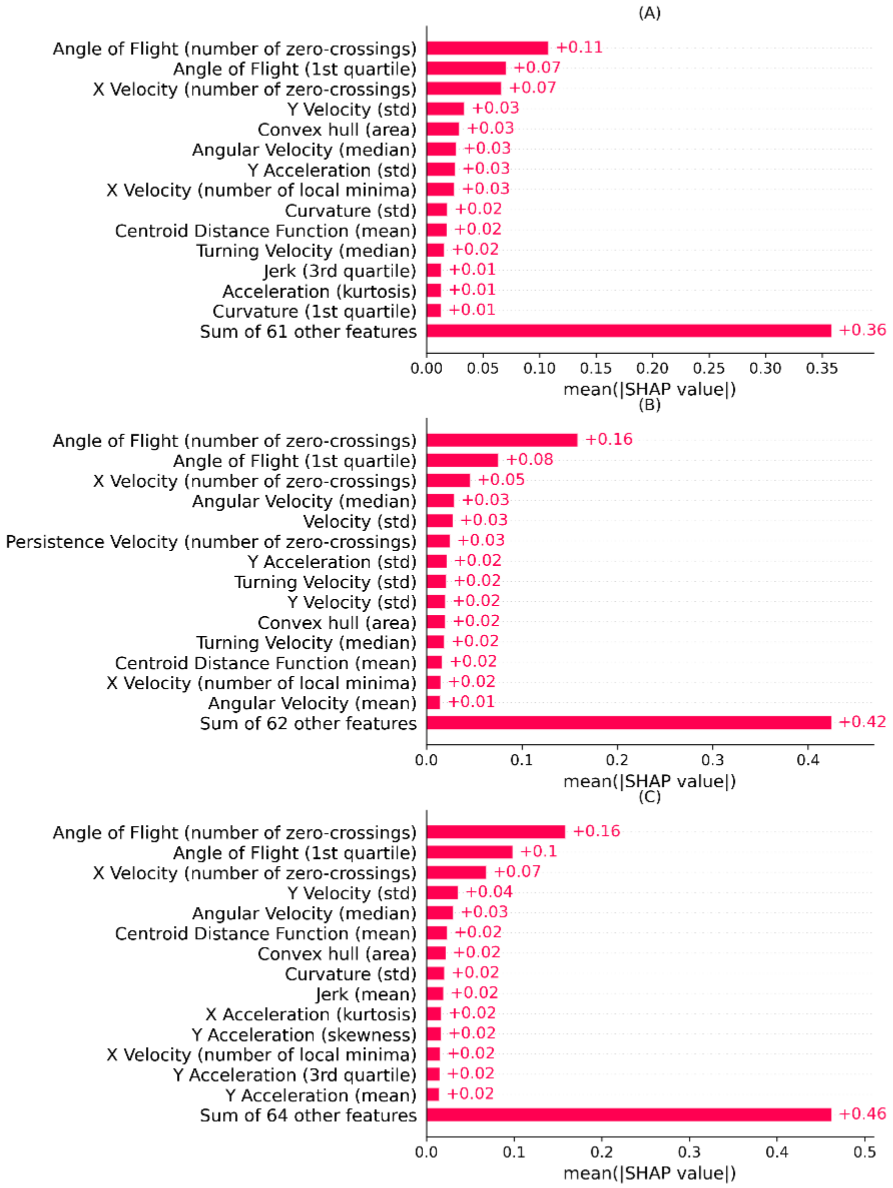


Figure 8. SHAP bar plots for the best model fold applied to independent test data for each data training strategy. In the figure, (A) displays the comprehensive data training strategy, (B) is the early data training strategy, and (C) is the balanced data training strategy. The features are ordered by mean absolute SHAP value.

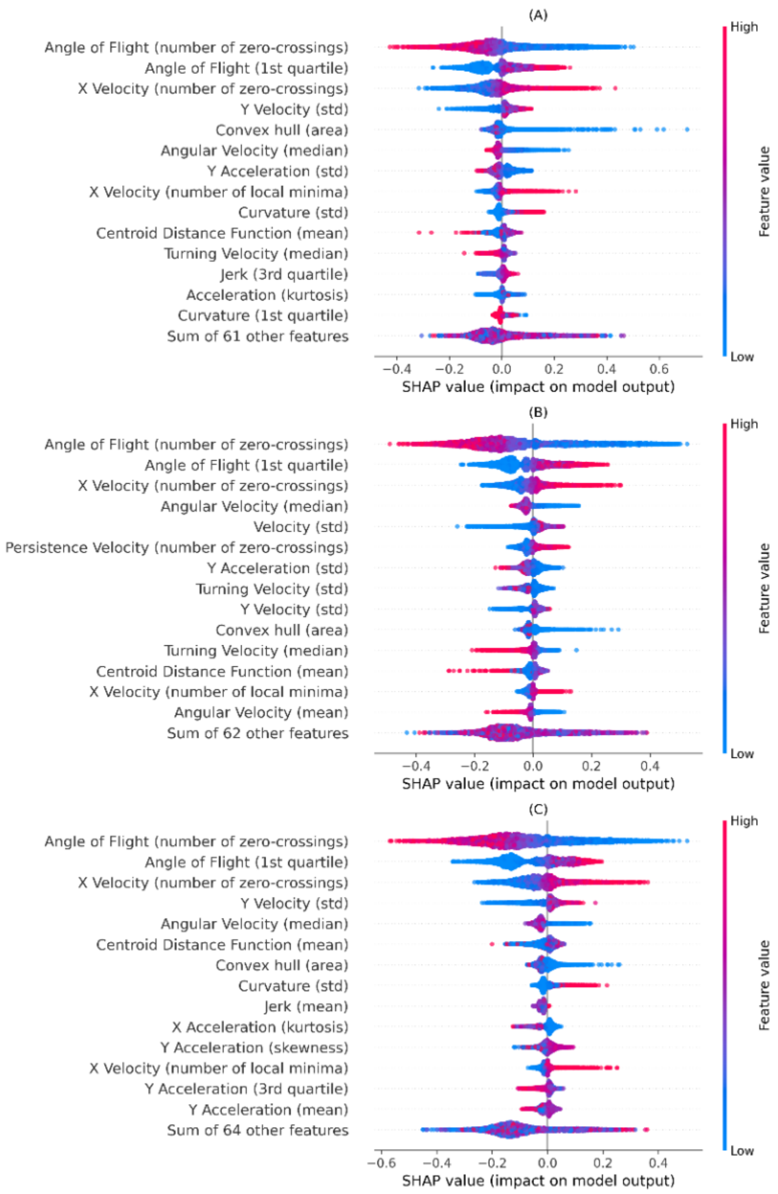


Figure 9. SHAP summary plots for the best model fold applied to independent test data for each data training strategy. Features are ordered by mean absolute SHAP value where each dot represents a segment, with its colour displaying its feature value. Positive SHAP values contribute towards the OL class, whilst negative SHAP values contribute towards the UT class. In the figure, (A) displays the comprehensive data training strategy, (B) is the early data training strategy, and (C) is the balanced data training strategy.

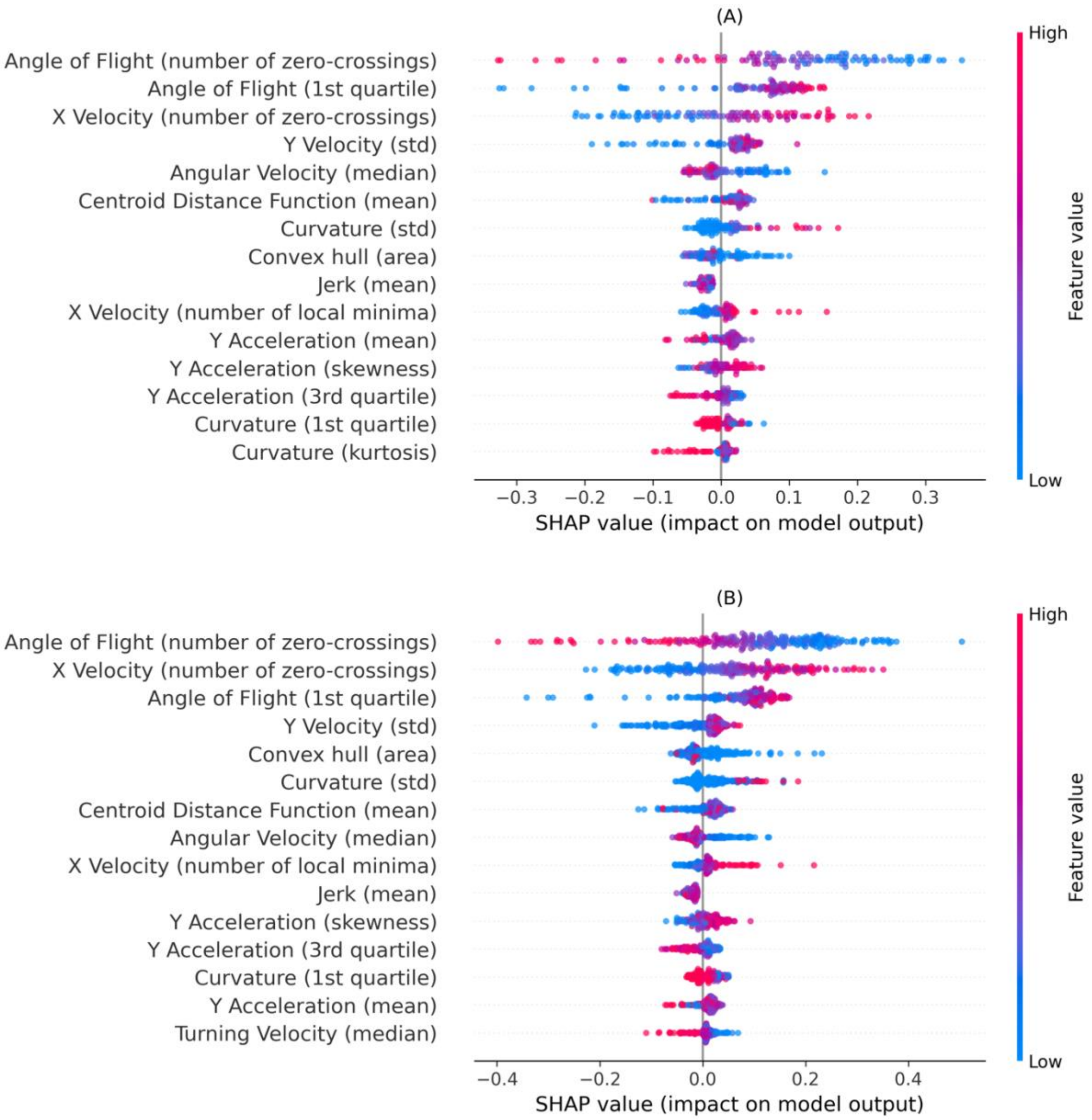


Figure 10. SHAP summary plots with IR and IS trajectories shown separately for the best model fold applied to independent test data using the balanced training strategy on the OL net. Features are ordered by mean absolute SHAP value where each dot represents a segment, with its colour displaying its feature value. Positive SHAP values contribute towards the OL class, whilst negative SHAP values contribute towards the UT class.  In the figure, (A) displays the IS class, and (B) displays the IR class.

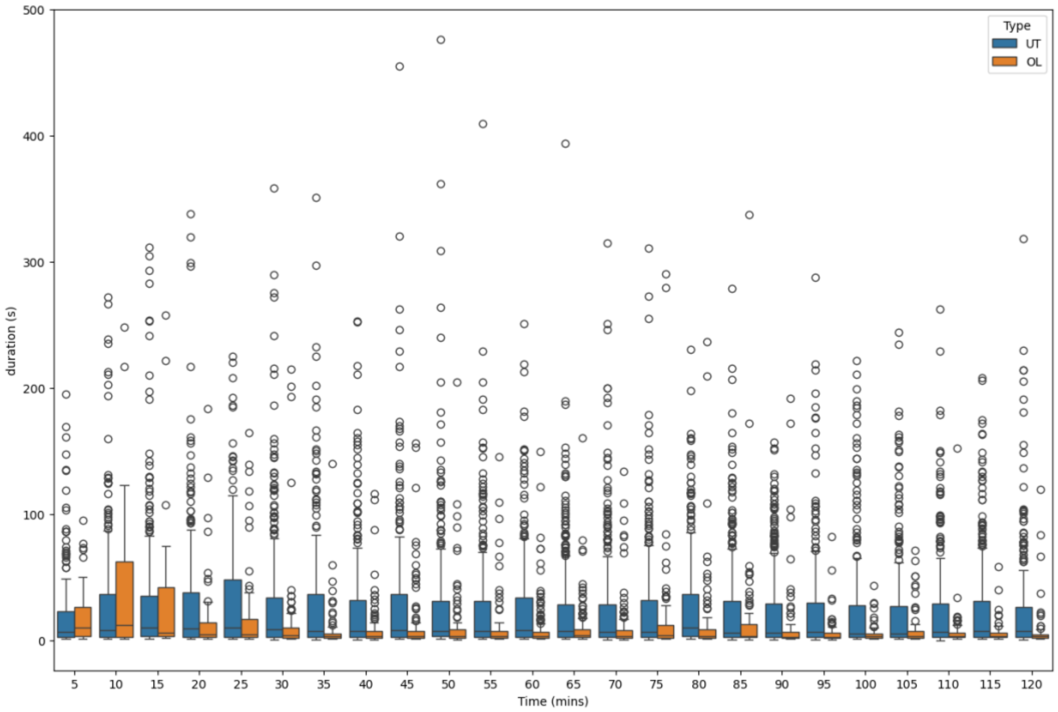


Figure 11. Boxplots of track duration for the Banfora strain across the time of the experiment. Both UT and OL datasets are displayed for comparison.

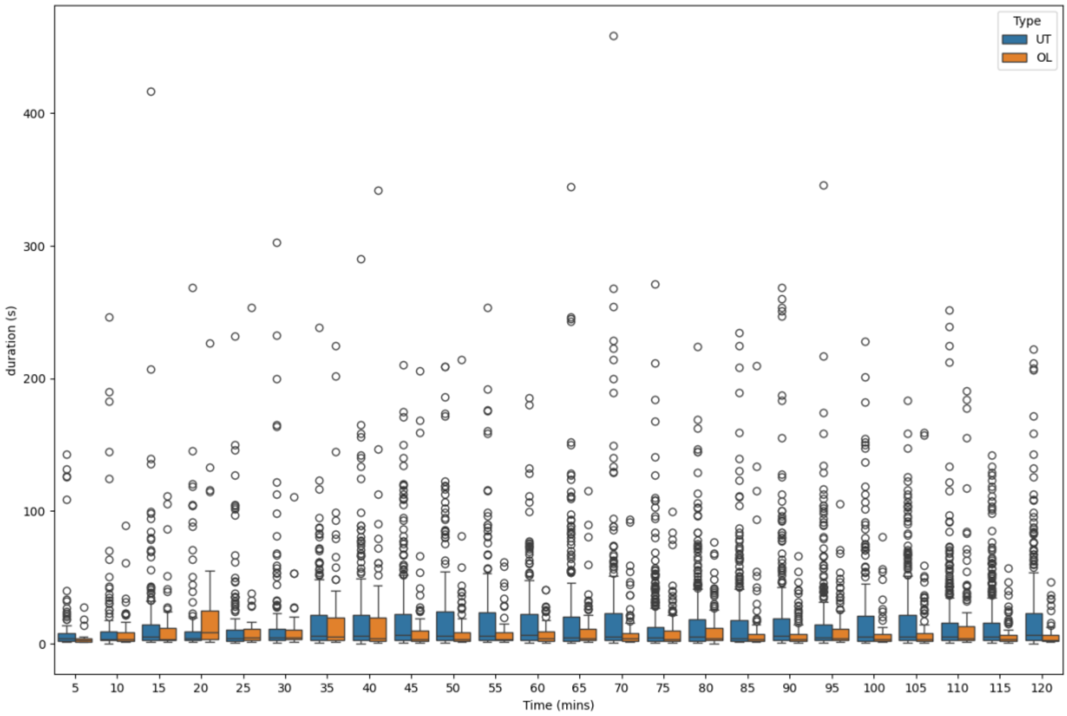


Figure 12. Boxplots of track duration for the VK7 strain across the time of the experiment. Both UT and OL datasets are displayed for comparison.

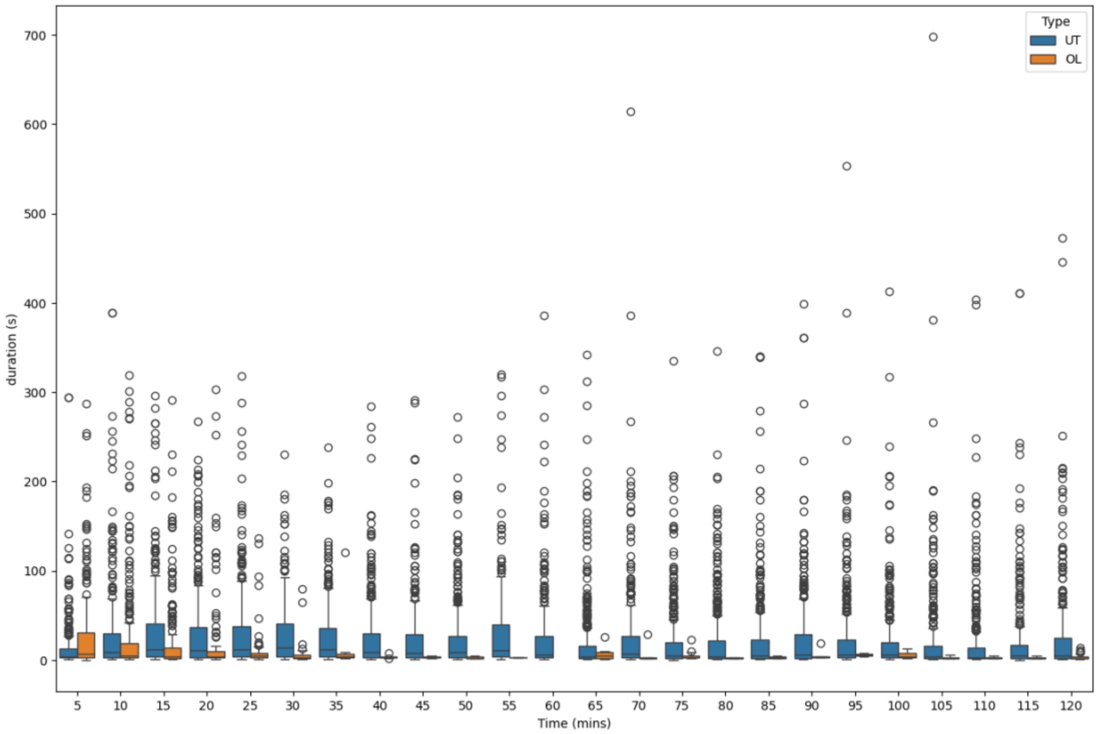


Figure 13. Boxplots of track duration for the Kisumu strain across the time of the experiment. Both UT and OL datasets are displayed for comparison.

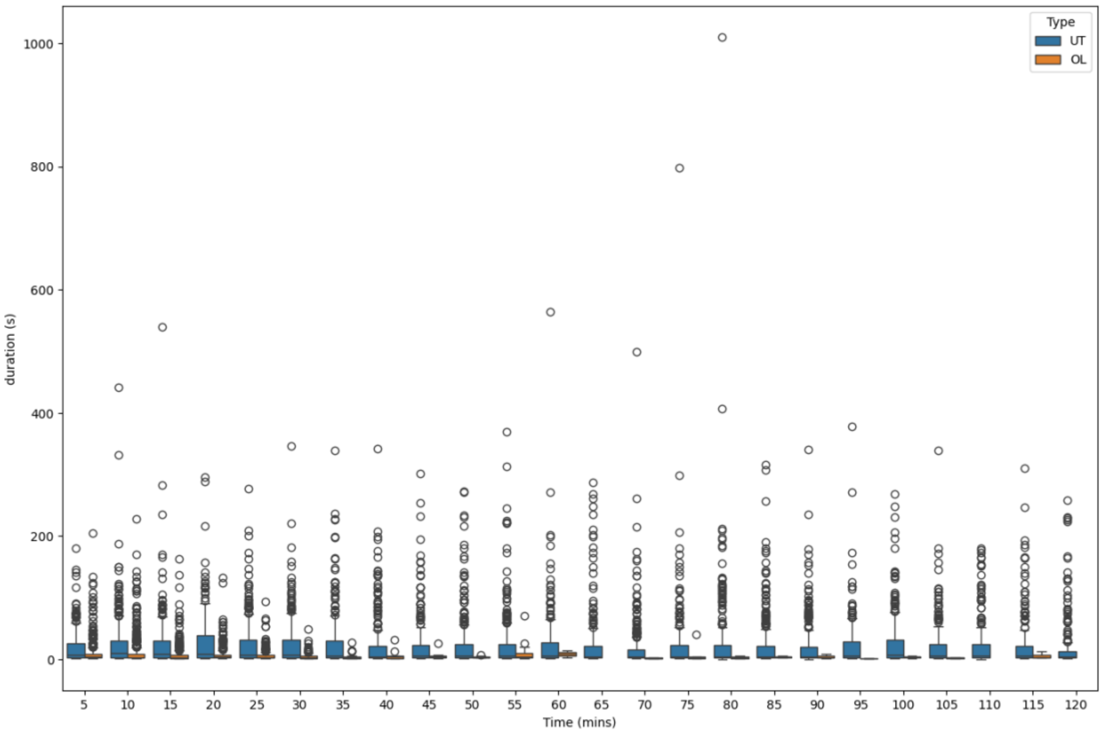


Figure 14. Boxplots of track duration for the Ngoussu strain across the time of the experiment. Both UT and OL datasets are displayed for comparison.

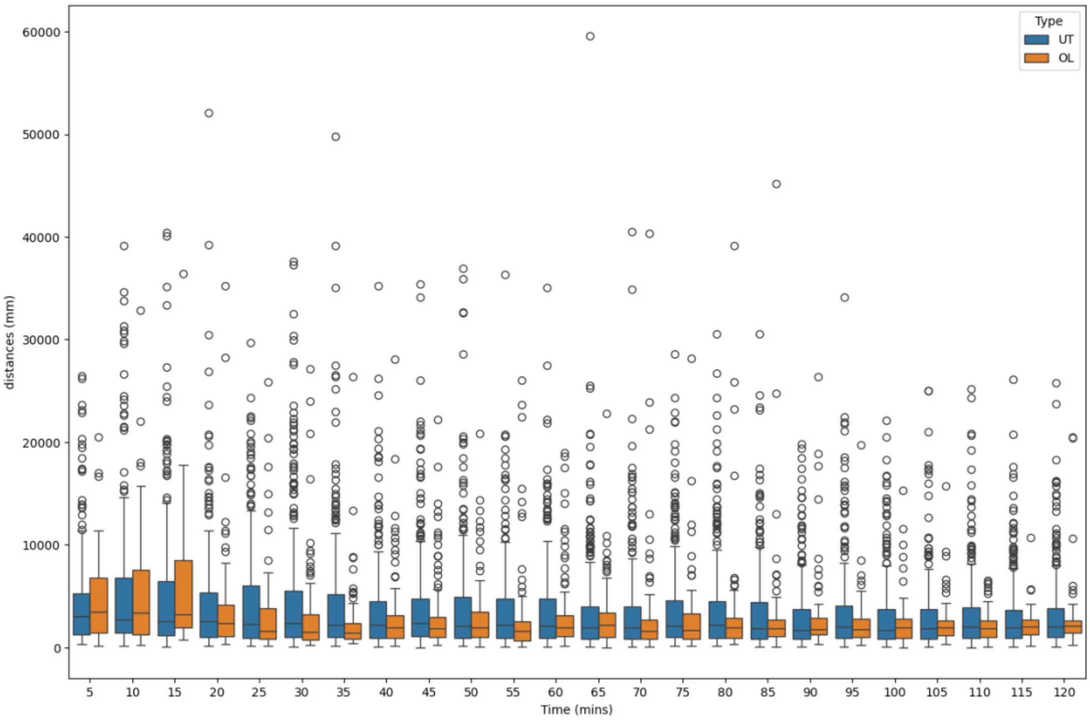


Figure 15. Boxplots of track displacement for the Banfora strain across the time of the experiment. Both UT and OL datasets are displayed for comparison.

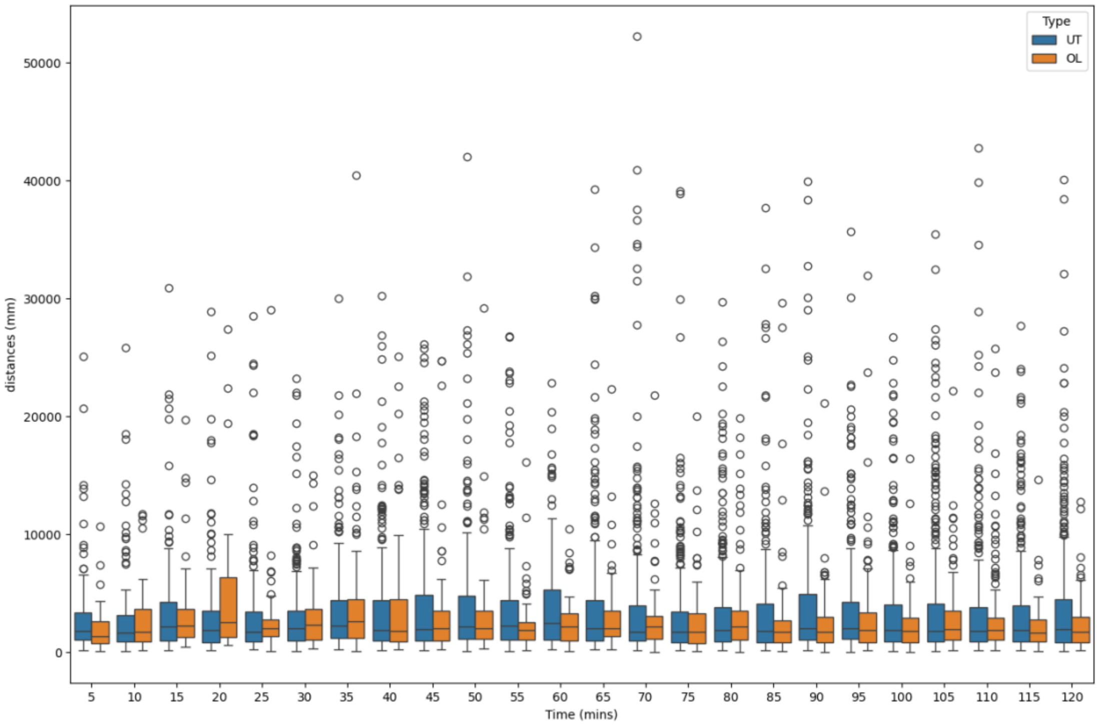


Figure 16. Boxplots of track displacement for the VK7 strain across the time of the experiment. Both UT and OL datasets are displayed for comparison.

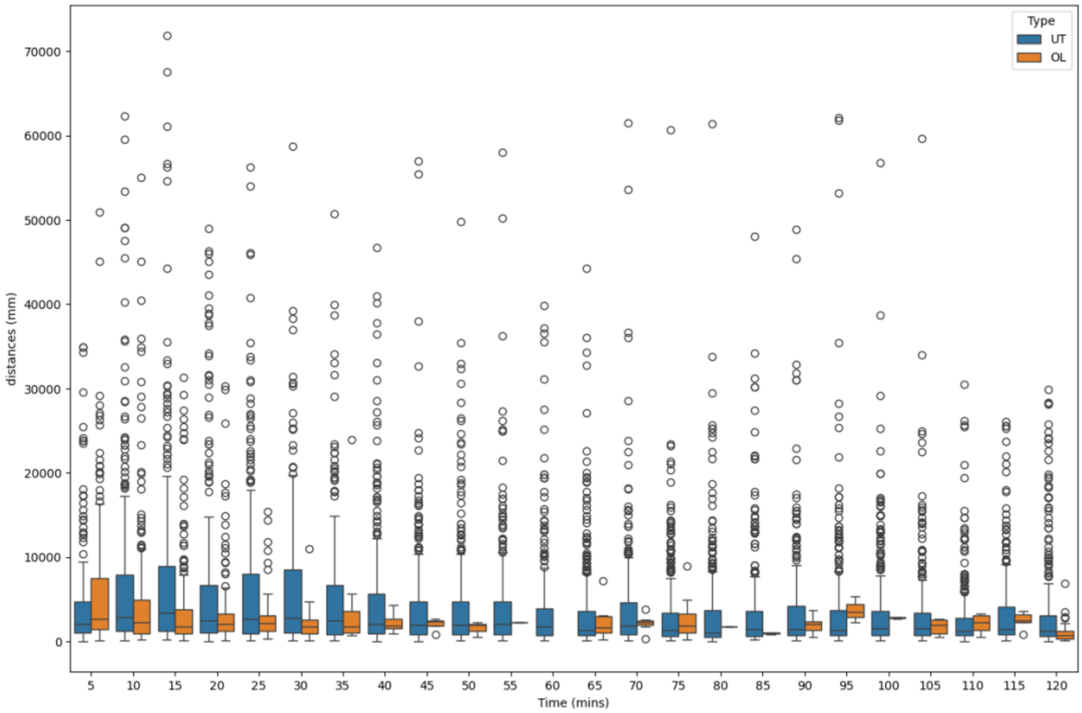


Figure 17. Boxplots of track displacement for the Kisumu strain across the time of the experiment. Both UT and OL datasets are displayed for comparison.

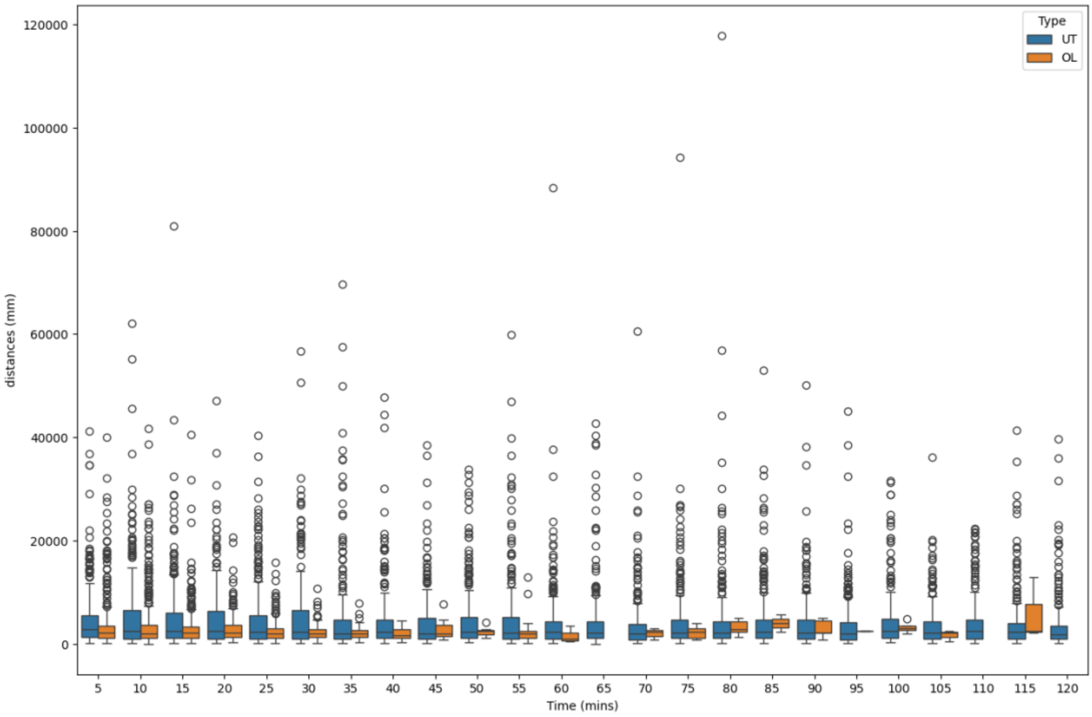


Figure 18. Boxplots of track displacement for the Ngoussu strain across the time of the experiment. Both UT and OL datasets are displayed for comparison.

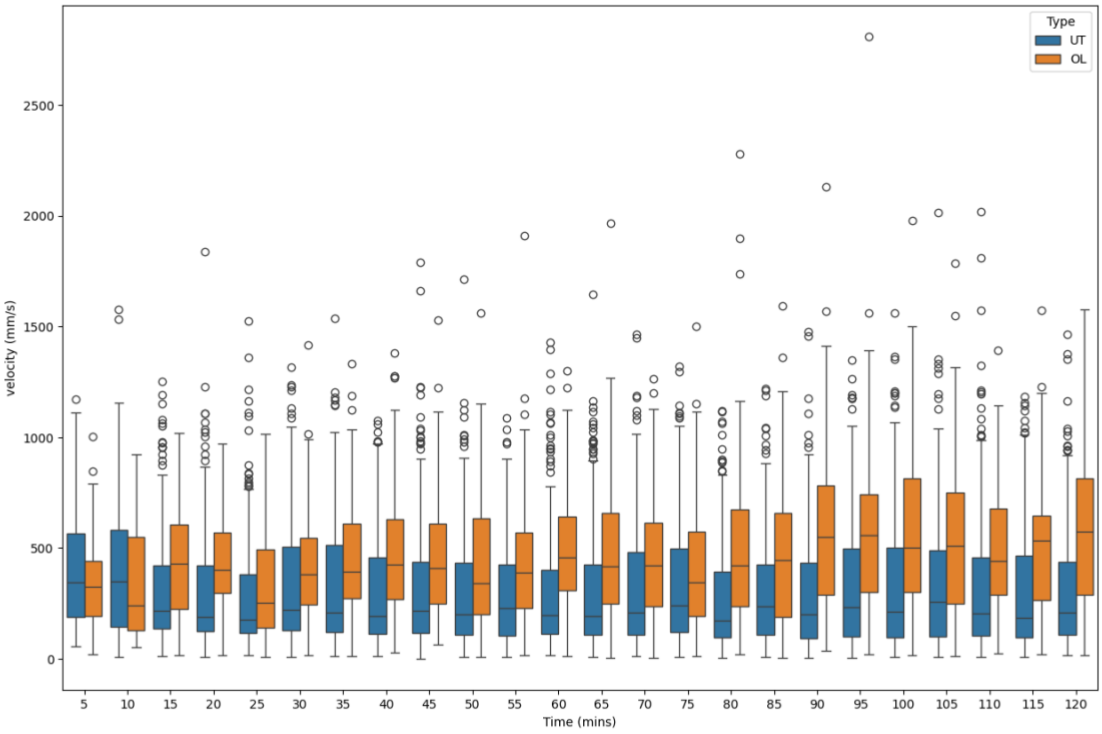


Figure 19. Boxplots of average track velocity for the Banfora strain across the time of the experiment. Both UT and OL datasets are displayed for comparison.

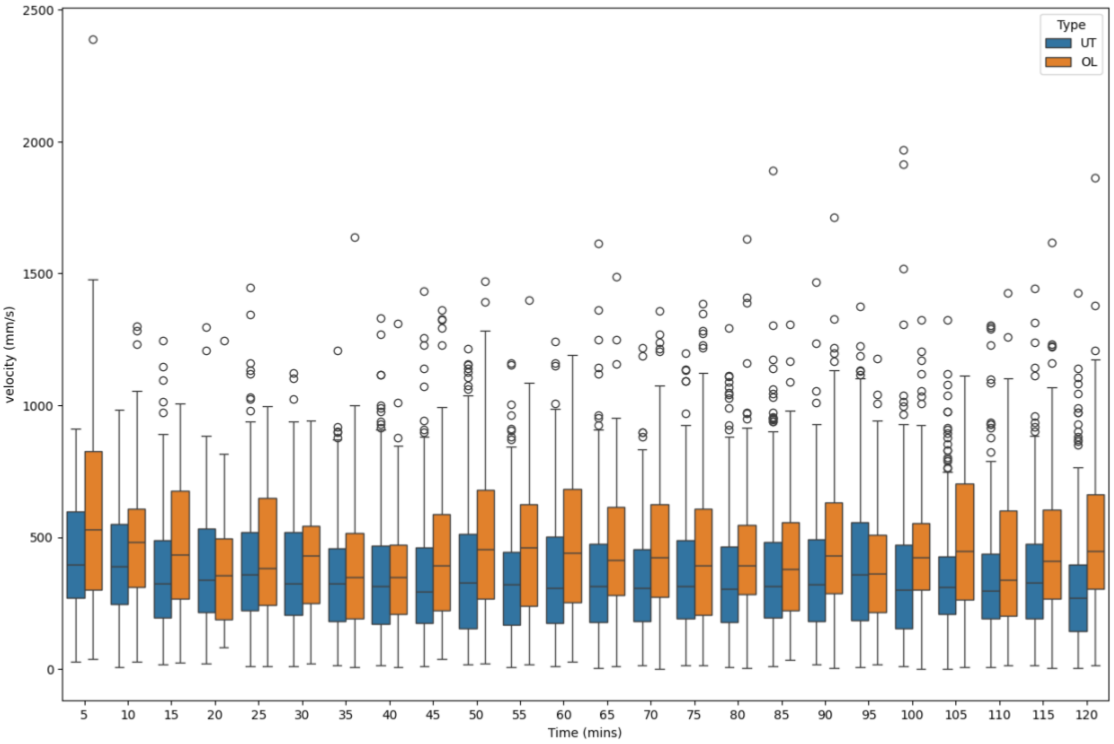


Figure 20. Boxplots of average track velocity for the VK7 strain across the time of the experiment. Both UT and OL datasets are displayed for comparison.

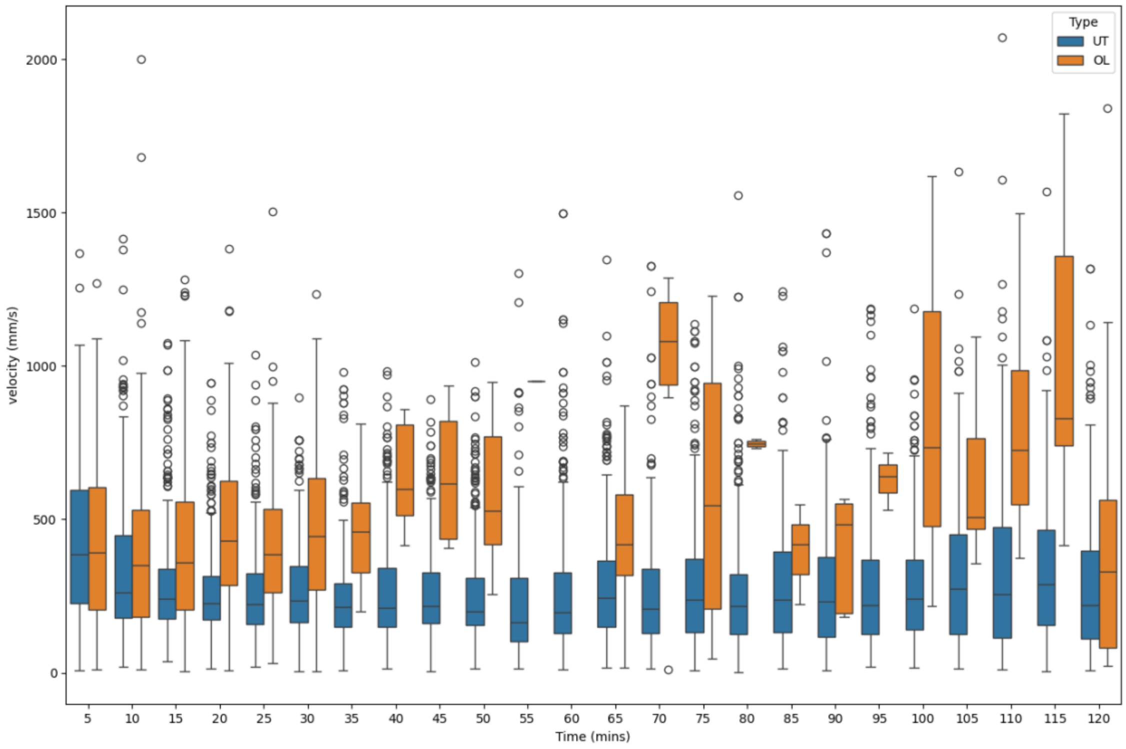


Figure 21. Boxplots of average track velocity for the Kisumu strain across the time of the experiment. Both UT and OL datasets are displayed for comparison.

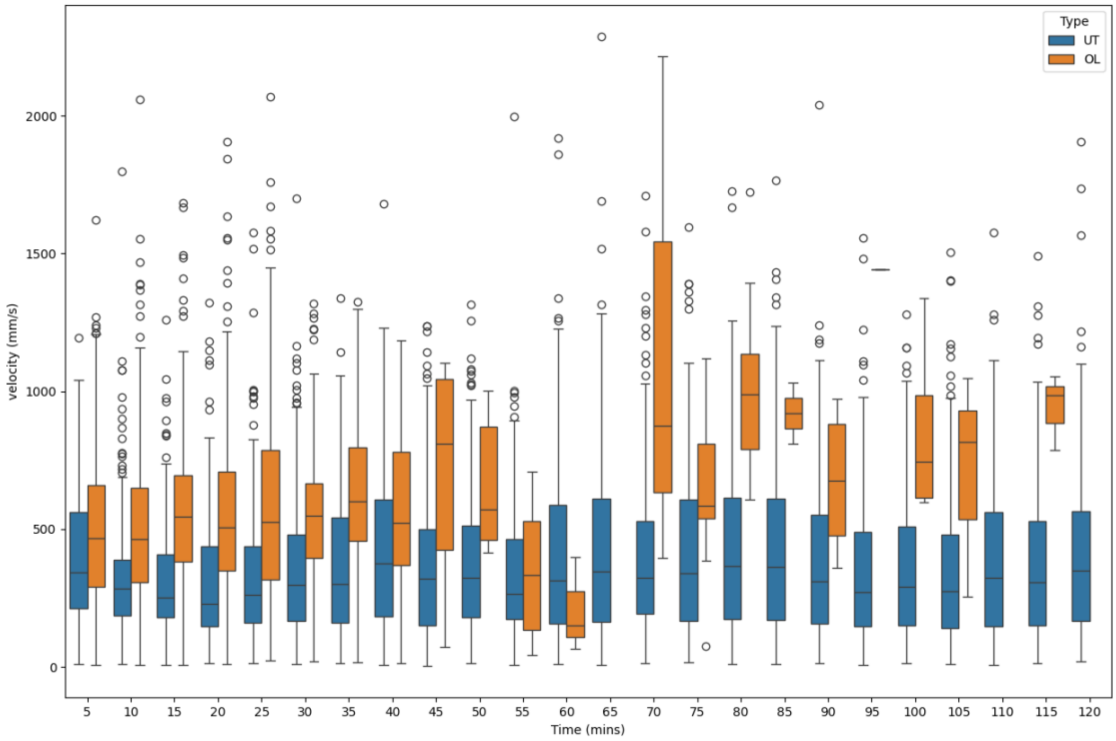


Figure 22. Boxplots of average track velocity for the Ngoussu strain across the time of the experiment. Both UT and OL datasets are displayed for comparison.
